# Identification of key host genes for influenza A virus in avian cells using a genome-wide CRISPR-Cas9 screen

**DOI:** 10.1101/2025.10.03.680283

**Authors:** Rosemary A Blake, Abraham Lee, Nicholas Parkinson, Spring Tan, Vayalena Drampa, Kenneth Bailie, Paul Digard, Mark Cigan, Rachel Hawken, Finn Grey

## Abstract

Influenza A virus (IAV) causes major economic losses to the poultry industry and poses a zoonotic threat to human health. Potential pandemic outbreaks are underpinned by the ability of the virus to jump from one species to another. Host-virus interactions can dictate the success of such events and while systematic studies have successfully mapped host virus interactions in human cells, few studies have been performed in relevant animal host cell lines. Here, we conducted two independent genome-wide CRISPR/Cas9 knockout screens in chicken lung epithelial cells infected with either the human-adapted PR8 vaccine strain or the avian UDL 3:5 reassortant virus encoding PR8 HA, NA and M segments. Rather than selecting solely for cell survival, we used anti-M2 antibody staining and fluorescence-activated cell sorting to capture host factors influencing multiple stages of the IAV life cycle. Across both screens, we identified 104 genes required for efficient replication in chicken cells, including 16 with strong effects (log₂ fold change > 2). Comparative analysis with published human screens revealed 17 conserved host factors, 19 human-specific factors, and 42 chicken-specific factors, highlighting potential species-specific interactions. Top hits included genes involved in sialic acid biosynthesis and N-linked glycosylation—*SLC35A1*, *SLC35A2*, and the avian-specific influenza polymerase cofactor *ANP32A*. Functional validation demonstrated that *MOGS*, *MGAT1*, *DENR*, *DMXL1*, *ENO1*, *IPO9*, *KLF6*, *PTAR1*, and *TSG101* contribute to multiple stages of the IAV life cycle. In particular, *MOGS* and *MGAT1* were essential for N-glycan processing and modulated cell-surface sialic acid abundance, with strain- and species-specific effects. These findings define a genetic landscape of IAV dependency factors in chicken cells and suggest shared and species-specific host requirements that could impact cross-species transmission.

## Introduction

IAV is a critically important pathogen for human health, food security and animal welfare. Despite coordinated international surveillance and control strategies, IAV regularly causes globally significant outbreaks of disease [1]. In humans, seasonal influenza outbreaks are a significant cause of morbidity and mortality, resulting in an estimated economic burden of $20 billion dollars annually in the US alone [2]. In terms of food security, poultry products are the major source of affordable animal protein throughout the world [3]. Production and consumption of poultry meat and eggs has increased markedly in the last 15 years [4]; in the same time span, IAV infections have resulted in the deaths of over 250 million birds [5, 6], whilst simultaneously raising plausible alarms over catastrophic human pandemics through regular zoonotic transmission events [7]. The current circulating highly pathogenic H5N1 subtype is evidence of this, and has been responsible for wild bird deaths [8, 9], increased mortality in poultry flocks [10], further expansion into new mammalian species [11–14] and some cases of human infection in recent years [7, 15].

Disease outcome from virus infection depends on complex host pathogen interactions. Rational design of therapies, vaccines and disease control depends on understanding these interactions at the molecular level. Due to resistance associated with targeting viral genes [16, 17], identification of key host-virus interactions provides important potential avenues for novel therapeutics. Furthermore, as viruses are obligate intracellular pathogens, they are completely dependent on the host cellular machinery for their replication. Precision editing of key host dependency factors (HDFs) represents a viable strategy for the generation of IAV-resistant farmed animals, as highlighted by recent studies on the avian specific HDF ANP32A [18, 19].

Genome-wide functional screens represent powerful approaches for systematic identification of host proteins involved in virus replication. RNAi screens and genome-wide CRISPR Cas9 screens have been extensively used to map human host interactions during influenza infection, identifying thousands of candidate host factors involved in IAV virus replication [20–29]. However, very few screens have been performed on cells from other species.

The pandemic nature of IAV is underpinned by the ability to jump from one species to another and historically every IAV pandemic has been sourced from an avian reservoir [30]. A lack of systematic screens in avian cells undermines our understanding of how influenza virus overcomes evolutionary barriers to jump from one species to another [31].

To address this gap in knowledge, we performed genome-wide CRISPR Cas9 loss-of-function screens in the chicken epithelial cell line CLEC213, infected with either lab-adapted PR8 (A/Puerto Rico/8/1934(H1N1)) or UDL 3:5 reassortant (A/chicken/Pakistan/UDL-01/2008 (H9N2)), an avian strain containing PR8 HA, NA and M segments. The level of M2 antibody binding to cells was used as a proxy of viral burden in each cell to assess the infection status of individual cells, in what we have termed a sort screen. The use of a sort screen allows the identification of a wide array of HDFs across the virus life-cycle, reducing the bias observed in survival screens which primarily allow the identification of host entry factors [32].

In this study, we identified factors previously identified in human screens, as well as novel hits previously not identified as associated with IAV replication. Detailed analysis of the two of the top hits, MOGS and MGAT1, demonstrated key roles in IAV infection of chicken cells. Contrary to earlier reports, MGAT1 is an important host factor in both chicken and human cells for IAV infection. This study establishes successful genome-wide sort screens in avian cells and identifies both common human and chicken HDFs while discovering new novel candidate chicken HDFs.

## Results

### Establishing genome-wide CRISPR Cas9 sorting screens in chicken cells

To identify avian host genes critical for IAV replication, we first generated a genome-wide CRISPR guide library cloned into the pKLV2-U6-gRNA5(BbsI)-PGKpuro2ABFP-W backbone. The chicken lentivirus library (GalGal6Cas9KO) encodes 70,484 sgRNAs targeting 17,233 protein coding genes (4 sgRNAs per gene), 754 miRNAs and 529 non-targeting sgRNAs as controls based on the GRCg6a reference genome. Amplification of the library in *E. coli* resulted in a 13.1 skew ratio and representation of 68594/71013 guides (96.6%), indicating a uniform representation of all the gRNAs in the library prior to transduction. To ensure efficient editing, we generated a clonal Cas9-expressing CLEC213 cell line (CLEC213-Cas9) by transducing wild-type (WT) CLEC213 cells with lentivirus expressing the Cas9 gene and tested individual clones for editing efficiency using a Cas9 reporter lentivirus (Supplementary Figure 1). We performed two independent pooled chicken genome-wide CRISPR screens challenged with PR8 and four screens with UDL 3:5 reassortant IAV (hereon in referred to as UDL) using a similar approach as described previously [20]. The use of PR8 allowed direct comparison to previously published human screens while the UDL virus allowed a greater understanding of infection of avian cells with an avian adapted strain. In brief CLEC213-Cas9 cells were transduced with the chicken genome-wide CRISPR library at an MOI of 0.3 to ensure the majority of cells represent single gene knock-outs (KOs), then selected using puromycin to eliminate non-transduced cells. On day 10-12 post transduction ∼300 million puromycin resistant cells were infected with PR8 or UDL at multiplicity of infection (MOI) of 5 for PR8 or 3 for UDL, achieving 95% infection based on M2 antibody fluorescent staining (Supplementary Figure 2). Cells were sorted by FACS into populations based on their levels of surface M2 expression (Figure 1A). Approximately 5% of cells were sorted into low or high populations and compared to a control population comprised of the mode of M2 expression (10-20% of the population).

**Figure 1.**
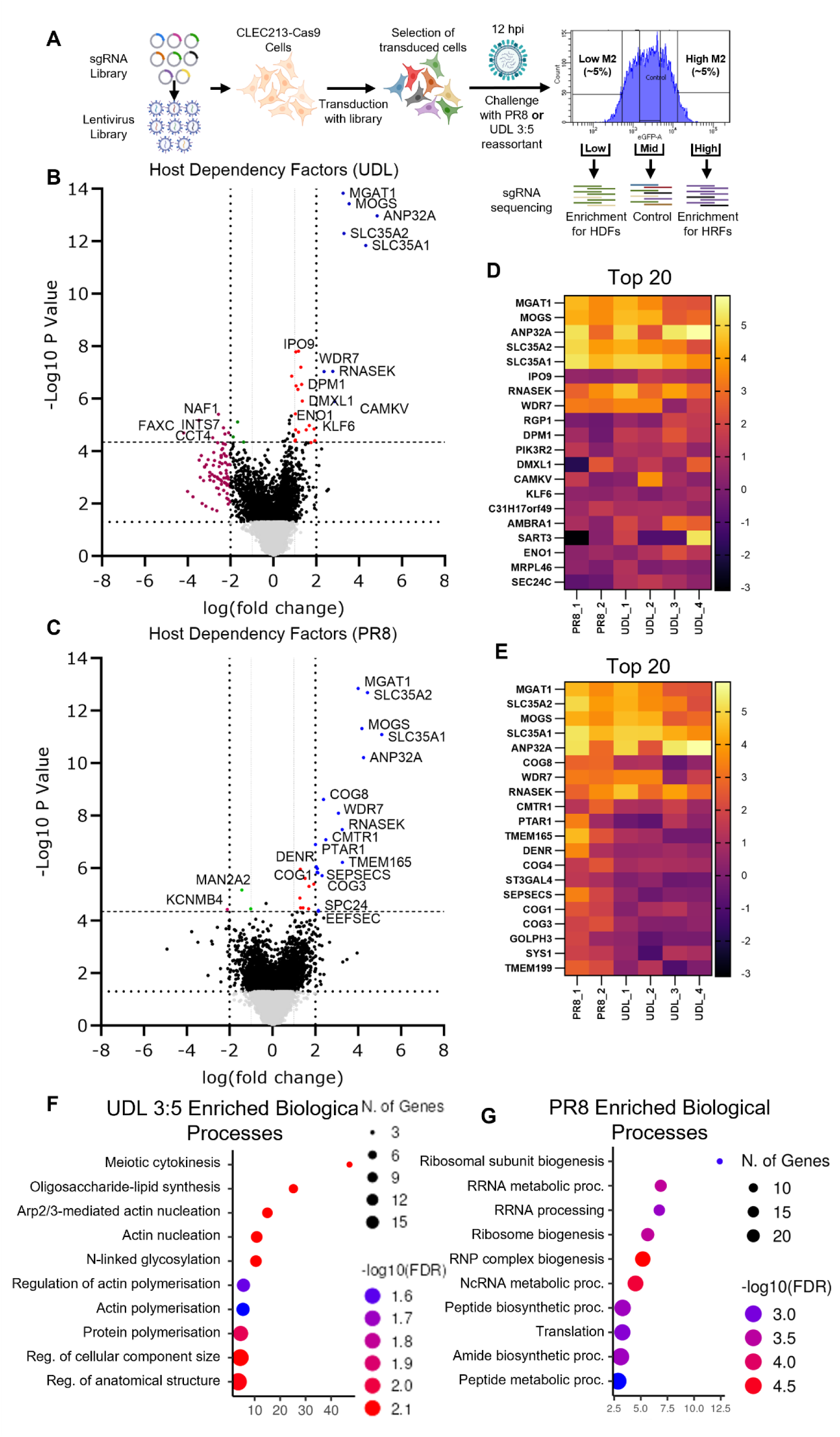
CRISPR sort screen to identify chicken host dependency factors (HDFs) in CLEC213 cells that promote IAV infection. (A) Schematics of the genome-wide CRISPR/Cas9 screening strategy. (B-C) MAGeCK analysis of multiple replicates comparing guides enriched in the “low” population to the “mid” population yielded log fold changes that were plotted on the *x*-axis and negative log10 P values that were plotted on the *y*-axis for both UDL (B) and PR8 (C) screens. (D) Heat map comparing the LFC of the top 20 HDFs between biological replicates in the UDL screen. (E) Heat map comparing the LFC of the top 20 HDFs between biological replicates in the PR8 screen. (F-G) Gene ontology analysis of UDL (F) and PR8 (G) HDFs (P<0.01).

### Identification of avian influenza virus host dependency factors

Integrated sgRNA cassettes were PCR amplified from each population of cells and submitted for Illumina Sequencing. Essential genes for chicken cells were analysed based on human genes (Supplementary Data 1) to analyse those which were considered universal essential genes from a range of human cell lines [33](long list) and a refined list of core essential genes based on network analysis of the long list [34](short list). Drop-out of these essential genes was observed over time in transduced cells under selection pressure, indicating successful editing (Supplementary Figure 3A). We observed robust coverage of the sgRNAs with an average of 358 guides per sgRNA for UDL and 586 guides per sgRNA for PR8 in in each sorted population (Supplementary Figure 3B). Candidate genes were identified and ranked using the model-based analysis of genome-wide CRISPR/Cas9 knockout (MAGeCK) program [35]. By comparing sgRNA representation in the low-M2 populations with that in the mid-M2 control populations, we identified 268 putative host dependency factors (HDFs) with *P* < 0.01 in the UDL screen (Supplementary Data 2) and 284 HDFs with *P* < 0.01 in the PR8 screen (Supplementary Data 3). Of these, 65 genes in the UDL dataset and 39 genes in the PR8 dataset met a false discovery rate (FDR) threshold of <0.25 (Supplementary Figure 3C).

In addition, known species-specific HDFs were identified, such as ANP32A, indicating the screens were successful in identifying host factors required by IAV to infect an avian cell. Comparing the Log Fold Change (LFC) against P value revealed eight HDFs with a LFC greater than 2 and an FDR <0.25 from the combined UDL screens (Figure 1B), and 16 HDFs with a LFC greater than 2 and an FDR <0.25 from the combined PR8 screens (Figure 1C). Of these, seven were enriched similarly in both screens and were ranked in the top 20 HDFs, five of which have been previously implicated in IAV infection; ANP32A, SLC35A1, SLC35A2, RNASEK and WDR7 (Figure 1 D-E). The LFC of the top 20 HDFs were observed to be similarly enriched between biological replicates of each viral screen and between viral screens, indicating a high level of reproducibility between biological replicates (Figure 1D-E).

Gene ontology (GO) analysis of genome-wide screens can inform our general understanding of cellular mechanisms that play important roles in IAV replication. Upon comparison of the HDFs, we identified 80 genes from the UDL and 163 genes from the PR8 screen mapped to known biological processes (*P*<0.01) (Figure 1F-G). We observed enrichment for HDFs involved in oligosaccharide-lipid synthesis and N-linked glycosylation, in addition to processes involved in RNA transcription and translation. These complexes have been similarly identified in human IAV screens and highlight the crucial roles in the viral-host interaction and subsequent intracellular replication of IAV [21].

### Comparative analysis of IAV screens in human cells

Previous genome-wide screens have identified multiple host factors important for IAV replication, predominantly from screens in human cells (Figure 2A). The pandemic nature of IAV is underpinned by the ability to jump from one species to another. However, such events require the virus to overcome barriers caused by evolutionary differences between host species. Understanding which host factors contribute to these barriers is essential for a full appreciation of the molecular processes that enable host jumping events and may improve surveillance and prediction of viral adaptations that increase pandemic risk. To assess conserved versus species-specific host requirements, we compared the hits identified in our chicken CRISPR screens with those reported in previously-published genome-wide IAV screens in human cells.

**Figure 2.**
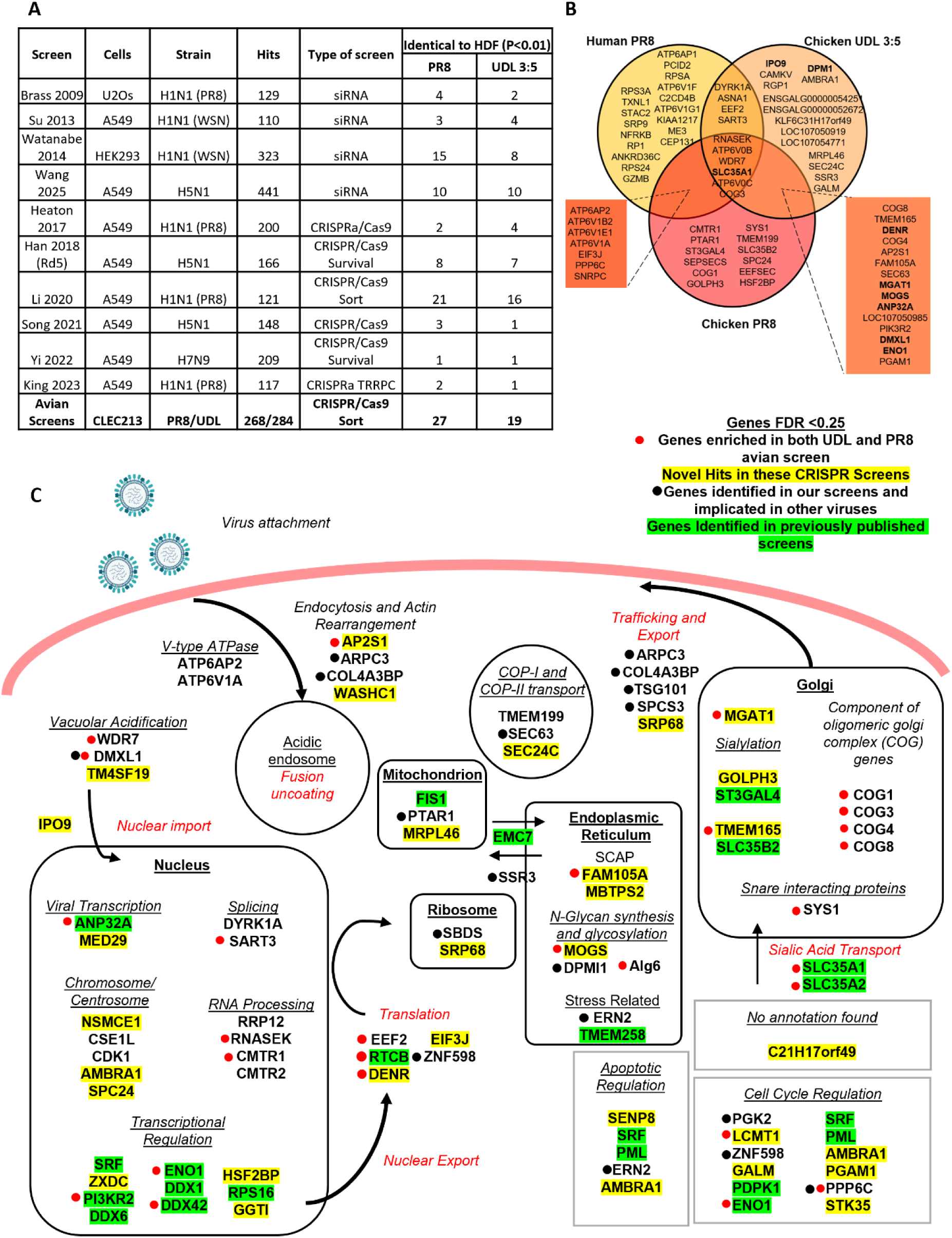
Comparison between avian and human HDFs against IAV. (A) A table comparing common HDFs from previously published genome-wide screens to HDFs *P*<0.01 from the UDL and PR8 avian screens. (B) Venn diagram comparing HDFs in the CLEC213 UDL, CLEC213 PR8 screen and A549 PR8 screen performed in the Baillie lab [20]. The top 30 ranked genes from each screen are included, with genes considered a hit in another screen if the *P* value was <0.05. (C) Avian CRISPR screen HDFs and their predicted roles in the IAV life cycle. Diagram shows all 104 HDFs from our CRISPR screens with both UDL and PR8 (FDR <0.25). Hits with a red dot were enriched in both UDL and PR8 sort screens. Hits unique to our CRISPR screens are highlighted in yellow. Hits with a black dot are those which have previously been associated with other viral infections prior to this study, but have not been associated with IAV. Hits highlighted in green are those which have been identified in previous human screens. All other hits are those enriched in only the UDL or the PR8 avian screen.

For comparability, HDFs with a *P*<0.01 were selected from the human and chicken sort screens for analysis with other screens to account for the differences in number of HDFs identified between screen types (Figure 2A). To this end, MAGeCK analysis was performed on the raw read counts from Li et al., 2020 human sort screen for more comparable analysis (Supplementary Data 4). 21 HDFs were shared between the chicken PR8 sort screen and the human PR8 sort screen, with 16 HDFs shared between the chicken UDL sort screen and the human PR8 sort screen (Figure 2A). All other genome-wide screens demonstrated shared HDFs with the chicken screens, including RNAi screens, but to a lesser extent (Supplementary Data 5). This both demonstrates the power of sort screens in their ability to identify a diverse array of HDFs, but also suggests that host species exerts diversity on identified HDFs and endpoint selection strongly affects screening outcome in addition to the IAV strain investigated.

The human CRISPR/Cas9 sort screen demonstrated the greatest number of shared HDFs and was therefore selected for further comparison [20]. To this end, the top 30 HDFs as ranked by MAGeCK analysis were selected to compare. This was to account for differences in the number of genes ranked as statistically significant by *P* value or FDR between the screens, whilst maintaining enough genes for a meaningful comparison to be made. Six HDFs were conserved between species and viruses (Figure 2B). These include SLC35A1, WDR7, COG3 and several V-ATPase subunits (Figure 2B). The enrichment of other V-ATPase subunits was particularly conserved for PR8 screens in both species. Unsurprisingly, there were a greater number of HDFs conserved between the two avian screens, indicating the identification of potential species specific HDFs. Of note, ANP32A was highly enriched in both avian screens and has been shown to be a species-specific HDF [18, 36] due to the redundancy between human ANP32A and ANP32B which both support IAV replication in human cells.

Combining data from the two sort screens conducted in this study revealed a total of 104 HDFs with FDR <0.25; 65 from the UDL screen and 39 from the PR8 screen. At least three sgRNAs were enriched for both screens for all genes shown and were evenly distributed across the chicken genome (Supplementary Figure 4). Figure 2C indicates the cellular role of HDFs identified from the screens and whether they are novel or have been previously associated with IAV replication or replication with other viruses. The prevalence of both characterised IAV HDFs in addition to novel hits indicates the robust data set generated from these avian screens and may indicate the possibility of species-specific HDFs. Certain genes such as DENR have not been implicated in any previous viral infection, and others such as DMXL1 have been characterised for other viruses [37] but have not been identified in previous IAV screens (Figure 2A, C) and therefore these may play an avian-specific role for IAV infection.

### Validation studies confirm top candidate hits from screens

Both MOGS and MGAT1 were highly enriched in the UDL and PR8 screens (Figure 1), but did not seem to be similarly enriched in human IAV screens or related previous research [38](Figure 2A-B) and were therefore selected for validation. Other genes selected for validation due to their enrichment in one or both avian screens and roles throughout the virus life-cycle include ANP32A, DENR, DMXL1, ENO1, IPO9, KLF6, PTAR1 and TSG101 (Figure 1A-B, 2C).

To validate selected HDFs, polyclonal KO cell lines were generated using the sgRNA which had the greatest level of enrichment in the “low” populations from the genome-wide screens. The polyclonal KO cells were infected at high and low MOIs with UDL or PR8 (Figure 3A-F). Upon confirmation of successful editing via sequencing, the polyclonal KO cells were infected with UDL (MOI 3) and PR8 (MOI 5). The level of M2 expression was determined 16 hours post infection (hpi) as this would replicate the readout of the CRISPR screen (Figure 3A-B). For both viruses, SLC35A1, ANP32A, MOGS and MGAT1 KO show greater than 50% reduction of M2 expression, and DENR and PTAR1 showed greater than 25% reduction in M2 expression (Figure 3A-B). Although all polyclonal KO cell lines created showed reduced M2 expression compared to the non-targeting guide control cells (NTG), DMXL1, ENO1, IPO9, KLF6 and TSG101 did not show significant reduction following infection with either UDL or PR8. While M2 expression levels provide a proxy readout for IAV replication levels, it is possible that some host factors may impact M2 expression without reducing virus replication. To provide a more direct assessment of the impact of candidate HDFs on IAV replication, viral titres were assessed by plaque assay 8 hpi (MOI 1) (Figure 3C-D) or 24 hpi (MOI 0.01) (Figure 3E-F) with either UDL or PR8.

**Figure 3.**
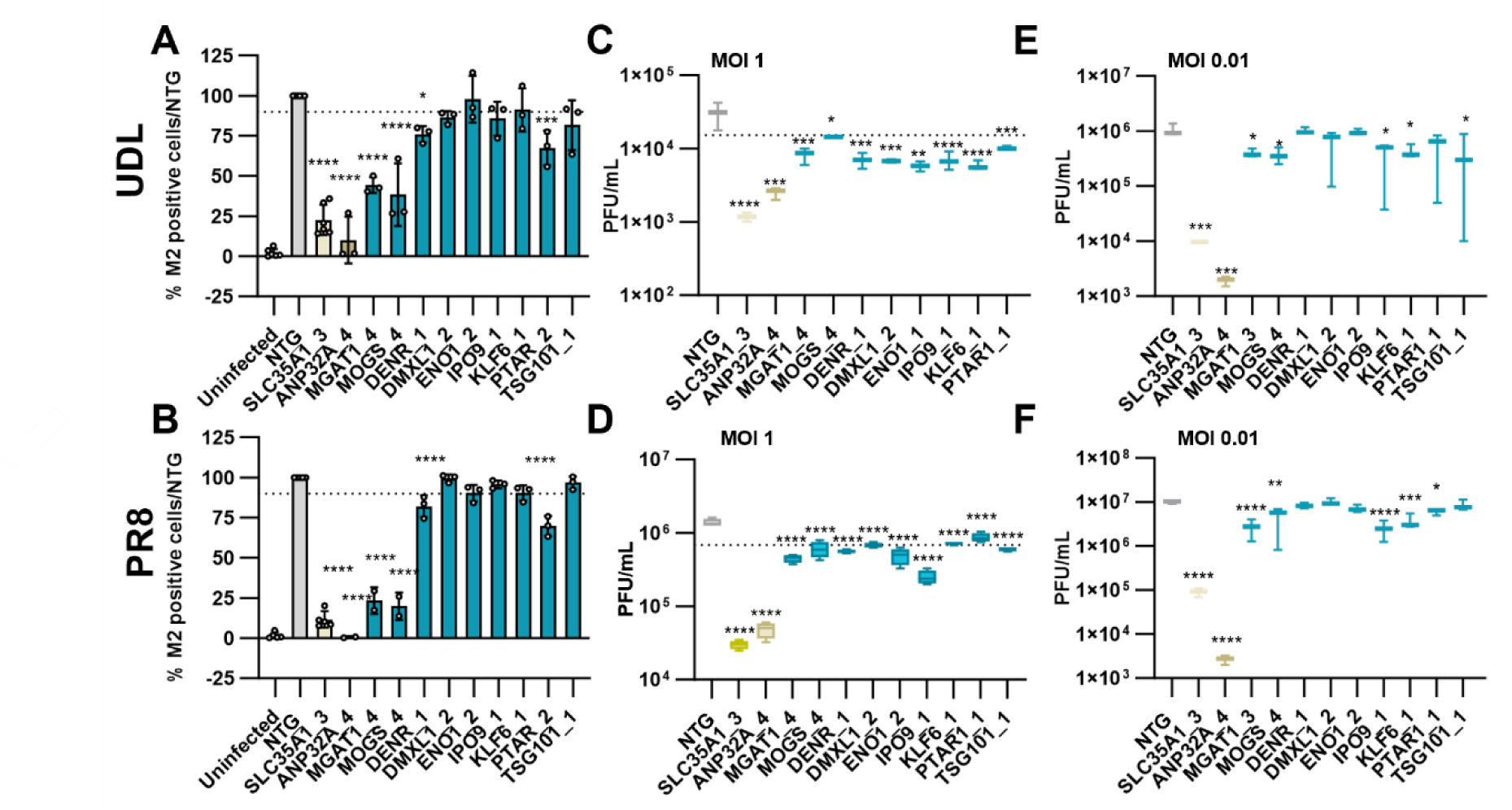
Validation of 11 candidate hits in CLEC213 cells with UDL 3:5 and PR8 IAV. CLEC213 polyclonal KO cells were transduced with either gene specific or non-targeting (NTG) sgRNA. Error bars represent standard deviations from three independent experimental replicates. CLEC213 KOs were infected with UDL or PR8 (A-F). (A-B) CLEC213s were infected with IAV at MOI 3 for 16 h. *Y*-axis shows percentage of M2-postive cells normalised to NTG. Dotted line denotes 10% reduction in M2 staining compared to NTG control. (C-D) Viral titre in plaque forming units (PFU)/mL at 8 hpi MOI 1 for UDL (C) or PR8 (D). Dotted line denotes ½ log reduction in viral titre compared to NTG for clarity. (E-F) Viral titre in PFU/mL at 24 hpi MOI 0.01 for UDL (E) or PR8 (F). *\*\*\*\*P*=0.0001 *\*\*\*P<*0.001 *\*\*P<*0.01 and *\*P*<0.05, by one-way ANOVA test followed by post-hoc Dunnett’s Test.

The viral titres for all HDF candidates were significantly reduced compared to the NTG control 8 hpi at high MOI for both UDL and PR8 (Figure 3C-D). At 24 hpi with MOI 0.01, seven of the ten candidates had significantly reduced viral titres. These were SLC35A1, ANP32A, MGAT1, MOGS, IPO9, KLF6 and TSG101 for the UDL (Figure 3E), and SLC35A1, ANP32A, MGAT1, MOGS, IPO9, KLF6 and PTAR1 for PR8 infected cells (Figure 3F). DENR and DMXL1 did not impact UDL or PR8 replication and TSG101 and PTAR1 acted in a strain dependent manner.

### Host factors involved in sialoglycan synthesis identified as key HDFs in chicken cells

IAV infection of a host cell is initiated by the binding of viral haemagglutinin (HA) to sialic acid (SA) moieties (sialoglycans) which are terminal sugars on glycans on the cell surface [21, 39, 40]. In both screens, enrichment of multiple HDFs involved in SA biosynthesis and the N-glycan processing pathway were observed (Figure 4A). MOGS (previously termed Glucosidase I [41]) is the first enzyme involved in the processing pathway for precursor N-glycans in the Endoplasmic Reticulum (ER), and MGAT1 (previously termed GntI [41]) is the enzyme that allows the conversion of high mannose N-glycans to hybrid or complex N-glycans, both of which have a SA moiety on the terminal glycan. Therefore, it was hypothesised these genes impact SA expression on the cell surface and may influence IAV infection.

**Figure 4.**
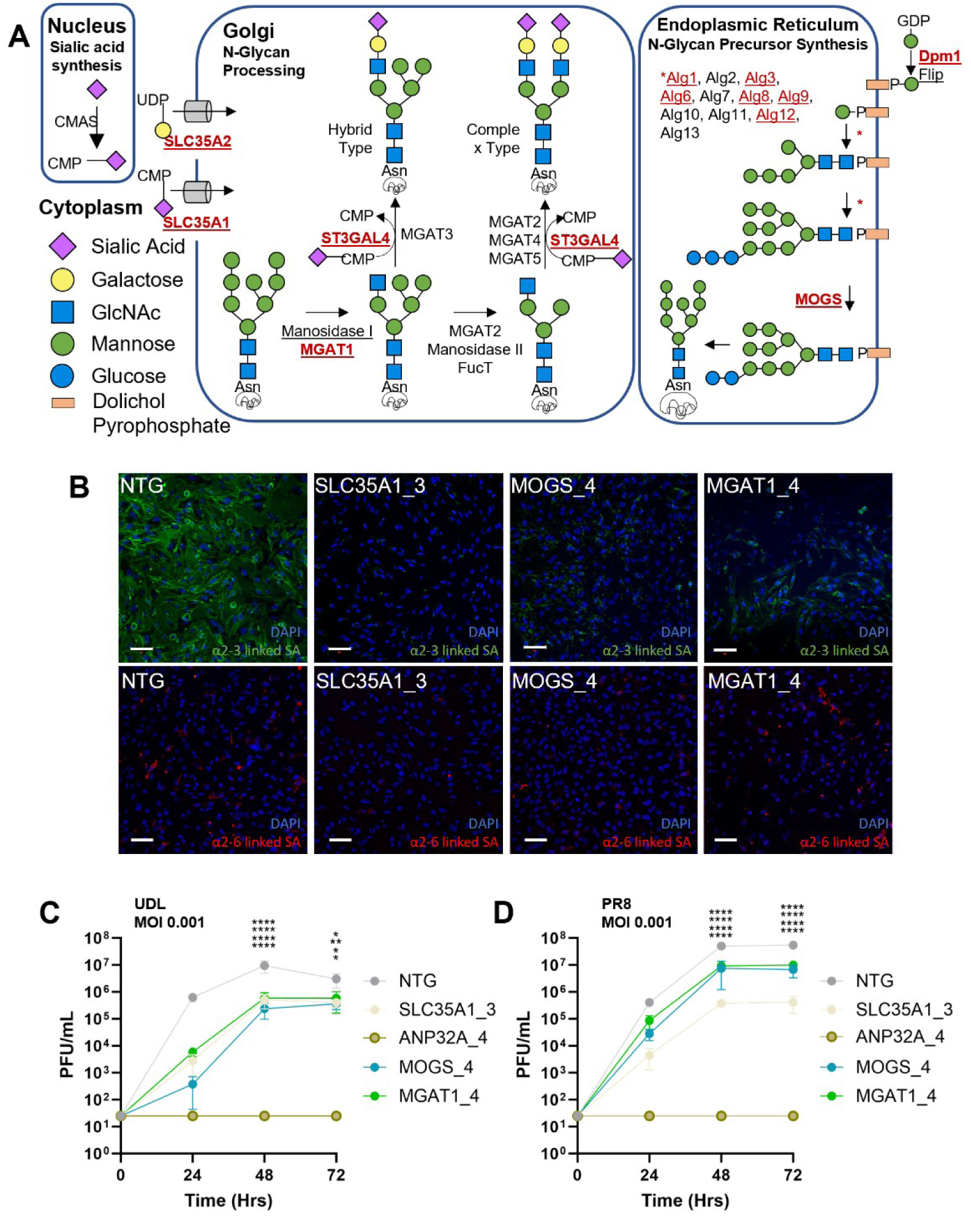
Function of MOGS and MGAT1 in CLEC213 cells. (A) Schematic of *de novo* sialic acid biosynthesis and N-glycan processing pathway. Significant genes identified in the GeCKO screens are shown in red. N-acetylglucosamine (GlcNAc), N-acetylmannosamine (ManNAc), cytidine monophosphate (CMP) and uridine diphosphate (UDP). (B) Staining for sialic acid surface expression with biotinylated antibody raised against MAL II (upper panel, green) or SNA (lower panel, red) lectin. Cells were counterstained with DAPI (nuclei, blue). (C-D) Virus titre in PFU/mL at 24, 48 and 72 hpi with either UDL (C) or PR8 (D) at MOI 0.001. Data representative of three biological repeats ±SD. *\*\*\*\*P*=0.0001 *\*\*\*P<*0.001 *\*\*P<*0.01 and *\*P*<0.05, by one-way ANOVA test followed by post-hoc Dunnett’s Test.

To assess this, NTG, SLC35A1, MOGS and MGAT1 KO cells were fixed and stained with lectin that binds to α2,3-linked SA (Maackia amurensis lectin, MAL II) or α2,6-linked SA (Sambucus nigra lectin, SNA) (Figure 4B). Confocal microscopy analysis showed control cells (NTG) containing a non-targeting guide, express α2,3-linked SA on the cell surface. However, expression was significantly reduced for SLC35A1, MOGS and MGAT1 KO cells. Little expression of α2,6-linked SA was observed for the CLEC213 cells.

Growth curves were conducted in the selected KO cell lines with UDL or PR8 at an MOI 0.001. Significant reductions in viral titres were observed in both KO cell lines, confirming their important role in IAV infection (Figure 4C-D).

### Phenotypic impact of N-glycan disruption depends on HA subtype

Different IAV strains with different HA proteins may have varying binding avidities to sialoglycans. This can depend on the virus binding preference for extended or shortened glycan receptors (H3N2)[42]; the ability of the virus to use alternative mechanisms of entry [43] or potentially the density of SA moieties linked to the glycan [38]. The type of HA expressed by IAV is known to affect the binding avidity of IAV to the cell surface, and while UDL and PR8 infection was reduced as measured by M2 staining and viral titre, both strains express the same H1 protein. To determine if avian IAV strains encoding different HA proteins have varying dependency on sialylated N-glycans, polyclonal MOGS and MGAT1 KO cell lines were generated using two separate guides from the genome-wide library, and infected at high and low MOI with three un-adapted low pathogenicity avian influenza strains: H5N2 (A/chicken/Belgium/150/1999), H6N1 (A/chicken/Netherlands/917/2010) and H9N2 (A/chicken/Saudi Arabia/2525/2000) (Figure 5A-I). For initial validation, the level of NP expression was determined 16 hpi with a high MOI (Figure 5A-C). For all strains tested, MOGS and MGAT1 polyclonal KO cell lines showed significant reduction in NP expression, with the exception of MGAT1_4 infected with H6N1. While significant, the reduction in viral staining for H5N2 and H6N1 was less than that observed for CLEC213 KOs infected with UDL (Figure 3A), PR8 (Figure 3B) and H9N2 (Figure 4C).

**Figure 5.**
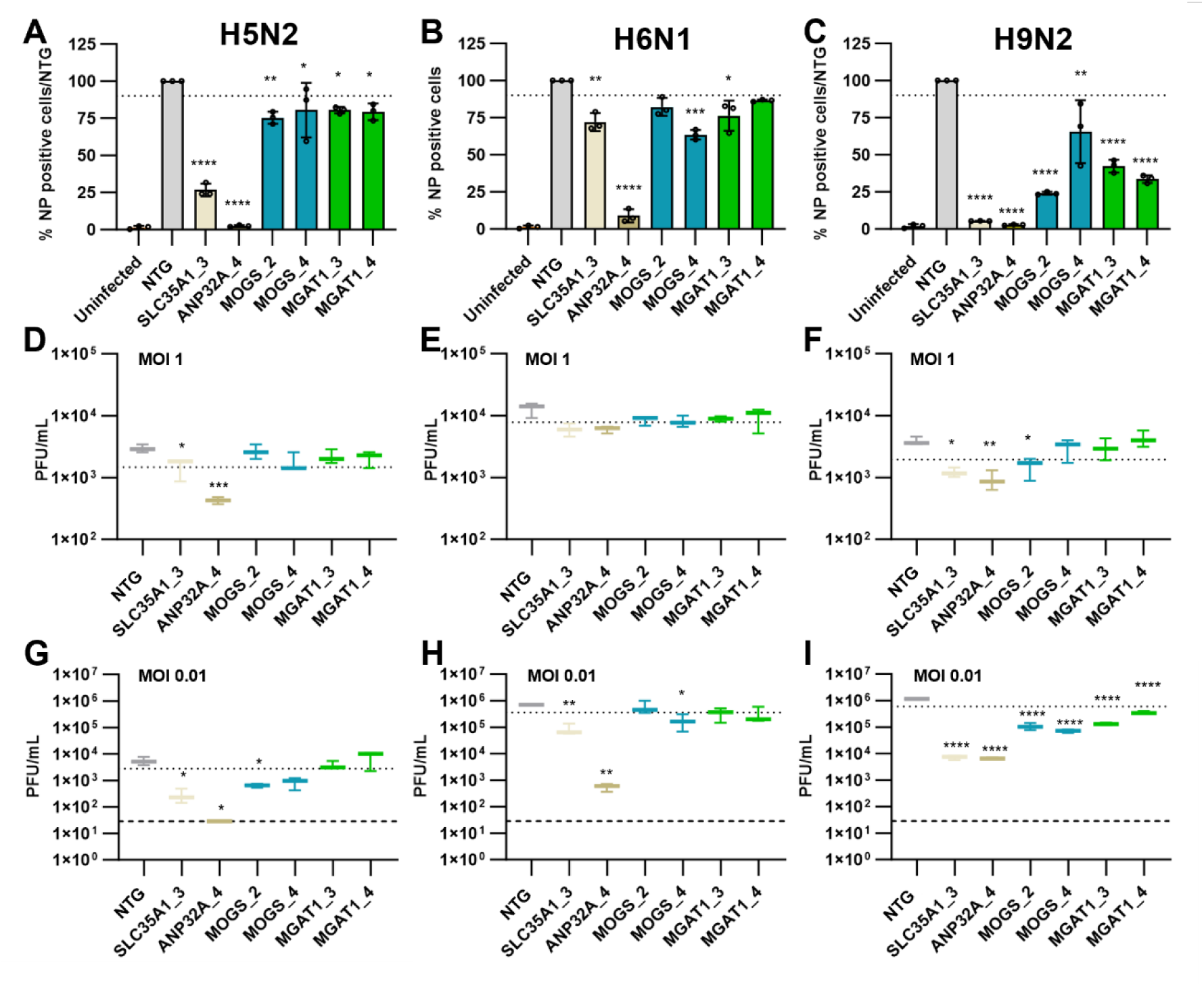
H5N2 and H6N1, but not H9N2, show robust replication in MOGS and MGAT1 KO CLEC213 cells. (A-C) CLEC213 cells with polyclonal MOGS and MGAT1 KOs infected with IAV at MOI 1 for 16 h. CLEC213 KOs were infected with (A) H5N2, (B) H6N1 or (C) H9N2 for 16 h. *Y*-axis shows percentage of NP^+^ cells normalised to NTG. Dotted line denotes 10% reduction in staining compared to NTG for clarity. (D-F) IAV titre in PFU/mL at MOI 1 8 hpi and (G-I) MOI 0.01 16 hpi for the indicated IAV strain. Dotted line for plaque assay data denotes ½ log reduction in viral titre compared to NTG. Dashed line indicates limit of detection for plaque assays. Error bars represent standard deviations from three independent experimental replicates. *\*\*\*\*P*=0.0001 *\*\*\*P<*0.001 *\*\*P<*0.01 and *\*P*<0.05, by one-way ANOVA test followed by post-hoc Dunnett’s Test.

When MOGS and MGAT1 KO cells were infected with MOI 1, there was no reduction in H5N2 (Figure 5D) or H6N1 viral titre (Figure 5E) at 8 hpi, and only one guide targeting MOGS had a significant reduction in H9N2 titre (Figure 5F). Furthermore, only one guide targeting MOGS resulted in a significant reduction in viral titre 24 hpi for both H5N2 (Figure 5G) and H6N1 (Figure 5H), while all guides targeting both MOGS and MGAT1 resulted in a significant reduction in H9N2 viral titre (Figure 5I) when infected with a MOI 0.01. The data suggests that MOGS and MGAT1 KO impact IAV infection, but the extent may be HA dependent, with greater reductions in viral infection observed for H1 and H9 IAV strains than H5 IAV strains (Figure 4-5).

### N-glycan expression impacting IAV infection is not cell-type specific

To establish if the impact of MOGS and MGAT1 KO on IAV infection was cell-type specific in chickens, MOGS and MGAT1 KO cell lines were generated using two separate guides from the genome-wide library in Cas9 expressing DF-1 cells. The cell lines were infected at high and low MOI with UDL and PR8 IAV (Figure 6). For initial validation, M2 expression was determined 16 hpi with MOI 3 (Figure 6A-B). A ∼10% reduction in M2^+^ cells for UDL infected cells (Figure 6A) and a significant reduction in M2^+^ cells for PR8 infected cells (Figure 6B) was observed compared to the NTG control, consistent with the reduction in M2^+^ CLEC213 KO cell lines observed previously (Figure 3A-B).

**Figure 6.**
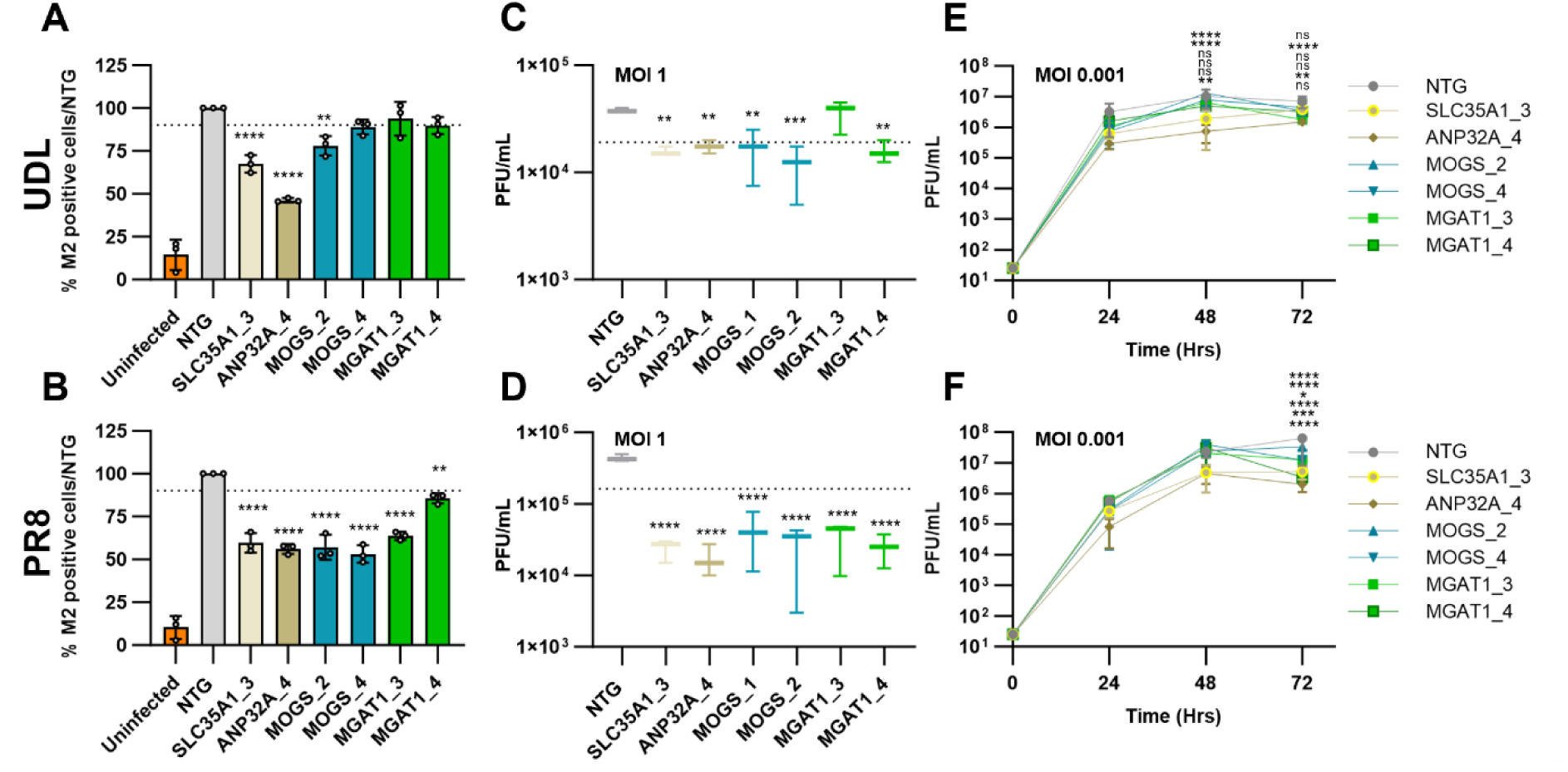
Validation of MOGS and MGAT1 in DF-1 cells. DF-1-Cas9 cells were transduced with either gene specific or NTG sgRNA. Two different sgRNAs were used to target MOGS and MGAT1 to generate individual polyclonal KO cell lines and were infected with UDL or PR8 IAV. (A-B) DF-1 cells were infected with IAV at MOI 3 for 16 h. *Y*-axis shows percentage of M2^+^ cells normalised to NTG. Dotted line denotes 10% reduction in M2 staining compared to NTG control for clarity. IAV titre in PFU/mL at (C-D) MOI 1 8 hpi (E-F) or at MOI 0.001 at 24, 48 and 72 hpi. Dotted line denotes ½ log reduction in viral titre compared to NTG for clarity. Error bars represent standard deviations from three independent experimental replicates. *\*\*\*\*P*=0.0001 *\*\*\*P<*0.001 *\*\*P<*0.01 and *\*P*<0.05, ns = non-significant, by one-way ANOVA test followed by post-hoc Dunnett’s Test.

When the polyclonal KO cells were infected with MOI 1, a significant reduction in viral titre was observed for UDL and PR8 8 hpi for the majority of cell lines investigated (Figure 6C-D). However, at 24 hpi (MOI 0.001) the majority of the polyclonal KO cell lines infected did not cause a significant reduction in viral titre for UDL or PR8 (Figure 6E-F).

It was concluded that while the reduction in IAV infection was not as great in DF-1 cell line compared to the equivalent CLEC213 cell line, MOGS and MGAT1 KO did impact IAV infection in both chicken cell lines suggesting the effect was not cell-type specific.

### N-glycan disruption impacts IAV infection in human cells

Depending on the chemical configuration, sialoglycans can act as a species-defined barrier to IAV infection. This is largely attributed to HAs from human IAV strains binding preferentially to α2-6 linked SA receptors and HAs from avian IAV strains binding preferentially to α2-3 linked SA receptors [44–47]. However, it is unknown if a specific type of glycoconjugate serves as the primary receptor across different species or if there are species variations. Previous research demonstrated that KO of MGAT1 has no impact on the ability of PR8 to infect A549 cells [38], and MOGS and MGAT1 were not identified as host dependency factors in any of the previously conducted human genome-wide screens with IAV (Figure 2). Furthermore, genes involved the processing pathways of other sialoglycans, such as O-glycans or glycolipids, were not enriched in the avian screens performed here (Figure 1B-C). Therefore, it was hypothesised that IAV infection of avian cells may be dependent on sialylated N-glycans over other SA-linked glycans, and this may be a species-specific phenotype.

To investigate if this phenotype was species-specific, A549 Cas9 cells were transduced with gene specific guides targeting MOGS and MGAT1. A significant reduction in M2 staining was observed in MOGS and MGAT1 polyclonal cell lines when infected with UDL or PR8 (MOI 3) (Figure 7A-B). MOGS and MGAT1 KO also resulted in reduced viral titres at 8 hpi for UDL infection (Figure 7C), but no significant differences were observed for PR8 infected cells 8 hpi (Figure 7D). Furthermore, differences in the viral titres from the cell lines for UDL or PR8 (MOI 0.001) infection were negligible (Figure 7E-F).

**Figure 7.**
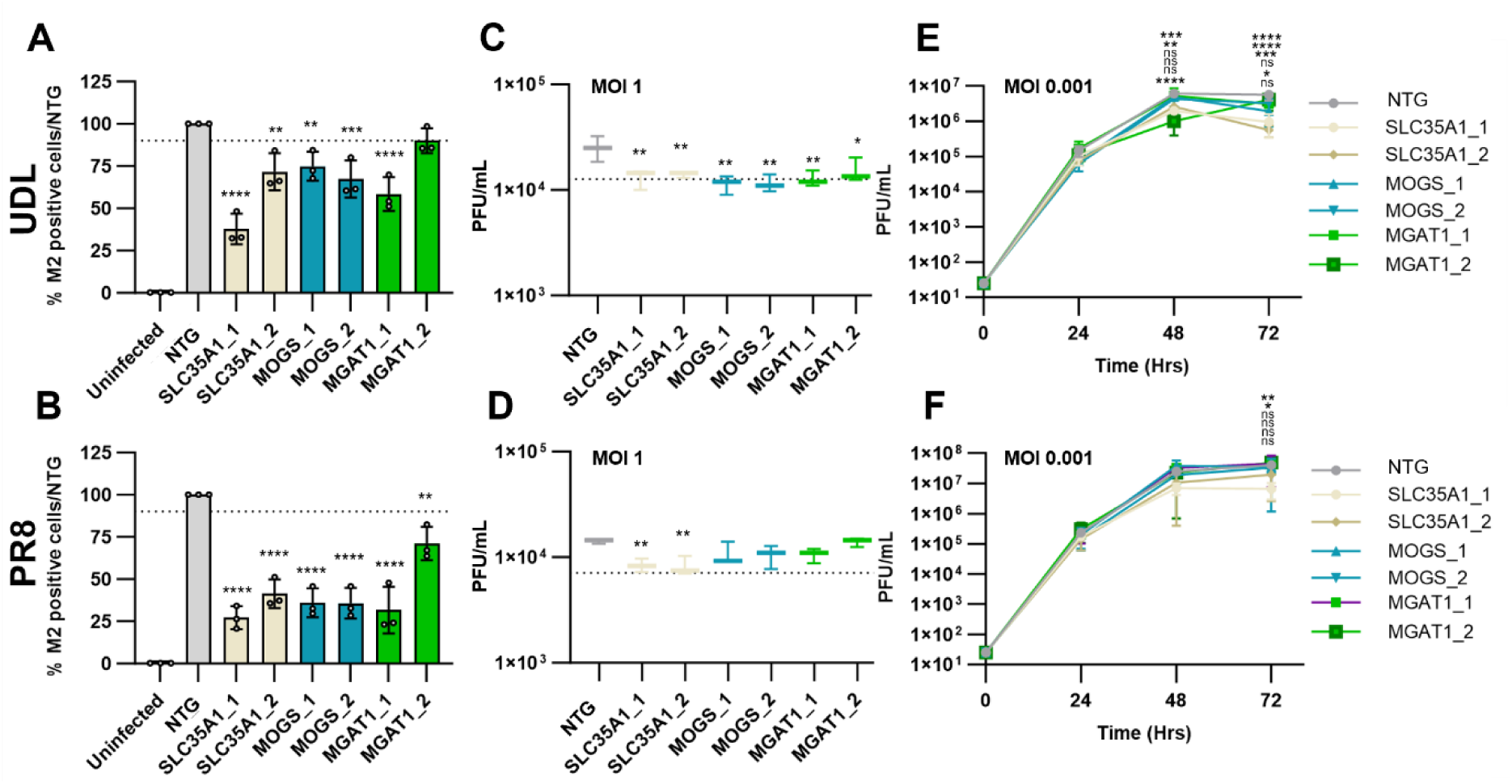
Validation of MOGS and MGAT1 in A549 cells. A549 cells were transduced with either gene specific or NTG sgRNA. Two different sgRNAs were used to target SLC35A1, MOGS and MGAT1 to generate individual polyclonal KO cell lines and were infected with UDL or PR8 IAV. (A-B) A549 cells were infected with IAV at MOI 3 for 16 h. *Y*-axis shows percentage of M2^+^ cells normalised to NTG. Dotted line denotes 10% reduction in M2 staining compared to NTG control for clarity. IAV titre in PFU/mL at (C-D) MOI 1 8 hpi (E-F) or at MOI 0.001 at 24, 48 and 72 hpi. Dotted line denotes ½ log reduction in viral titre compared to NTG for clarity. Error bars represent standard deviations from three independent experimental replicates. *\*\*\*\*P*=0.0001 *\*\*\*P<*0.001 *\*\*P<*0.01 and *\*P*<0.05, ns = non-significant, by one-way ANOVA test followed by post-hoc Dunnett’s Test.

While the reduction in IAV infection was less severe in DF-1 and A549 KO cells compared to CLEC213 MOGS and MGAT1 KOs, the attenuated phenotype was also observed for the viral titres harvested from the SLC35A1 KO in DF-1 and A549 cells (Figure 6E-F, 7E-F). This may be attributed to the reduced Cas9 editing efficiency in DF-1 Cas9 and A549 Cas9 cells (Supplementary Figure 5), and the remaining unedited protein in these cell lines is sufficient to produce viral titres equivalent to that of the NTGs over time.

## Discussion

In this study, genome-wide CRISPR/Cas9 sort screens in avian cells were performed, using both the avian-adapted UDL and the murine-adapted human PR8 IAV strains. This dual-strain approach enabled direct comparison of host factor requirements across viruses with different adaptation histories, providing new insights into IAV–host interactions in chickens. While previous work has applied survival-based genome-wide CRISPR screens in chicken DF-1 fibroblasts [48], the use of an antibody-based phenotypic read-out at early time points allowed identification of HDFs acting across multiple stages of the viral life cycle [20].

Multiple factors previously implicated in IAV infection, including multiple ATPases (*ATP6AP1*, *ATP6V0C*, *ATP6V0D1*) and known entry factors (*SLC35A1*, *WDR7*, COG-complex genes) were identified [20, 21, 49]. In addition, both UDL and PR8 screens identified *ANP32A*, an avian-specific cofactor for the viral polymerase [50], confirming identification of species-specific HDFs alongside broadly conserved ones.

HDFs absent from earlier genome-wide analyses in either human or chicken cells were also identified, thereby adding to current understanding of IAV–host interactions in avian systems. 27 novel HDFs were identified across the IAV replication pathway, 7 of which were enriched in both avian screens (*MOGS, MGAT1, DENR, TMEM165, AP2S1, FAM105A, LCMT1*) and a further 13 HDFs which were not implicated in previous human IAV research but have been demonstrated to have a role in the host-pathogen interaction of other viruses *(DMXL1* [37]*, DPM1* [51]*, TSG101* [52], *PTAR1* [53], *ARPC3* [54]*, COL4A3BP, SPCS3* [55]*, SSR3* [56]*, SBDS* [57]*, ZNF598* [58]*, ERN2, PGK2, PPP6C* [59].

Validation experiments confirmed 11 HDFs from the screens as important for replication of both UDL and PR8 in chicken cells, with most acting within a single replication cycle. DENR, DMXL1, and ENO1 impaired infection at early but not later stages, suggesting stage-specific roles. Of particular interest were *MOGS* and *MGAT1*, top-ranked hits in both screens and absent from human CRISPR datasets. These enzymes act sequentially in N-glycan processing, which may influence the presentation of sialic acid (SA) receptors used by IAV for attachment [41].

Our data indicate that N-glycan–linked SA presentation is important for IAV infection in chicken cells, with *MOGS* and *MGAT1* knockouts significantly reducing infection by multiple subtypes (H1N1, H9N2) and to a lesser extent by H5N2 and H6N1. This pattern suggests subtype-specific requirements for SA receptor density. Notably, *MOGS* and *MGAT1* knockouts in human A549 cells caused significant reduction in infection for both UDL and PR8, contrary to previous reports [38]. These results imply that while N-glycan–linked SA is not the sole determinant of IAV infection in human cells, its contribution is conserved across species.

Finally, while this study focused on HDFs, comparing “high” versus “mid” M2⁺ cell populations allowed identification of host restriction factors (Supplementary Figure 6). Several candidates emerged, including DCPS, DPH1, CHAF1B, and SRSF11, with DCPS and SRSF11 significantly enriched in both screens. Given their roles in mRNA de-capping and splicing—processes essential for IAV replication—these HRFs are plausible antiviral effectors and warrant further investigation [60, 61]. However, few classical antiviral genes were identified, such as interferon (IFN) related genes, suggesting this is not the optimal approach for identifying HRFs or redundancy obscures their identification. Potential future experiments could include pre-treatment of cells with IFN prior to infection with IAV, or infection with IAV mutants unable to counter the IFN response.

In conclusion, our study demonstrates the utility of genome-wide CRISPR/Cas9 sort screens in avian cells, enabling the identification of both conserved and species-specific HDFs and HRFs. The discovery of novel factors such as *MOGS* and *MGAT1* expands our understanding of glycan–virus interactions and provides potential targets for genetic approaches to enhance influenza resistance in poultry. These findings also highlight the value of cross-species comparative screens in revealing fundamental aspects of viral biology and host specificity.

## Methods

### Cell culture and viruses

Chicken lung epithelial cells (CLEC213 cells), a kind gift from Dr Sascha Trapp, French National Institute for Agriculture, Food and Environment, France) [62] and chicken fibroblast cells (DF-1 cells) were cultured in DMEM/F12 (Gibco) supplemented with 10% foetal bovine serum (FBS; Gibco), 1% penicillin/streptomycin/glutamine (PSG; Gibco) and 15 mM Hepes buffer. Human embryonic kidney cells (HEK293T cells), human lung carcinoma cells (A549 cells), Madin-Darby canine kidney cells (MDCK cells) and MDCK cells overexpressing chicken ANP23A (MDCK_chANP32A^+^; donated by Prof Massimo Pamarini, University of Glasgow Centre for Virus Research, UK) [63] were cultured in DMEM (Gibco) supplemented with 10% FBS and 1% PSG. The DF-1-Cas9 and CLEC213-Cas9 cell lines were generated from transduction with lentivirus generated from lentiCas9-Blast (Addgene #52962) and selected for using 10 µg/mL blasticidin (Invivogen). The A549-Cas9 cell line was kindly provided by Professor Kenneth Baillie [20]. PR8 (A/Puerto Rico/8/1934(H1N1) was generated through reverse genetics in our laboratory and propagated in MDCK cells [64]. UDL 3:5 reassortant (A/chicken/Pakistan/UDL-01/2008 (H9N2), a kind gift from Prof Wendy Barclay, contained all the internal segments of the UDL H9N2 virus with the HA and NA segments from PR8 to mitigate safety concerns and the M segment to allow screening with 14C2), H5N2 (A/chicken/Belgium/150/1999), H6N1 (A/chicken/Netherlands/917/2010) and H9N2 (A/chicken/Saudi Arabia/2525/2000) strains were kindly provided by Sjaak de Wit and propagated in eggs and MDCK cells in our laboratory [65].

### Chicken library generation

A genome-wide CRISPR-Cas9 knockout (GeCKO) sgRNA library was designed based on *Gallus gallus* genome assembly GRCg6a, using custom Python scripts. Gene annotations were combined from Ensembl release 99 and Refseq release 104, favouring genes with assigned symbols or longer coding sequences where there were conflicting annotations between datasets at a given locus. For each gene, designs were based on the transcript with the greatest exon use concordance compared to other transcripts. For protein-coding genes, candidate sgRNAs were identified based on presence of an NGG protospacer-adjacent motif (PAM) and Cas9 cut site within an exon within the first 5-65% of the coding sequence. Designs were also included for miRNAs where possible, with Cas9 cut sites within the stem-loop sequence. After removing candidates containing a ‘TTTT’ U6 stop codon, *BSmbI* restriction site or homopolymer sequence > 5 bases in length, on-target scores were calculated using Rule Set 2 scores [66]. To screen for potential off-target effects, all possible matches to the genome were identified with up to 3 mismatches, with either an NGG PAM or one of the lower efficiency PAMs NAG, NCG, NGA, NGC, NTG or NGT, and an off-target score was computed by calculating the Cutting Frequency Determinant score [66] for each match and aggregating scores using the formula described by Hsu et al. [67]. A high impact off-target score was additionally calculated for all off-target matches targeting exons or non-coding RNA genes. To minimise effects of genetic variation, guides were screened against chicken variant data from Ensembl release 99 and the CFD score calculated to model the impact of common variants on cutting efficiency. Final selection of sgRNAs was based on a composite score derived from on-target and off-target scores, position in the gene, frequency of target exon usage and lack of impact of variation, prioritising sgRNAs spaced apart by more than 5% of the coding sequence. Finally, non-targeting control sgRNAs were identified by screening candidate 20mers against the genome as for the targeting guides, and selecting sgRNAs with negligible predicted off-target effects. The final designed library contained 70,484 targeting sgRNAs targeting 17,987 genes (17,233 protein-coding genes and 754 miRNAs) and 529 non-targeting controls. For 596 genes, unique targeting was impossible due to sequence identity with closely-related genes, and so sgRNAs were shared with paralogues for joint targeting. A mean of 3.9 sgRNAs per gene were selected (range 1-4). The designed oligonucleotide pool was synthesised and cloned into the pKLV2-U6gRNA5(BbsI)-PGKpuro2ABFP-W vector by a commercial service provider (Azenta Life Sciences, Burlington, MA, USA) and amplified in Stb3 *E. coli*. The library was purified and transfected with with pAX.2 (Addgene #12260) and pMD2.G (Addgene #12259) packaging plasmids and lipofectamine 2000 to generate the lentivirus library using HEK293T cells.

### CLEC213 genome-wide CRISPR screen and data analysis

CLEC213 cells were transduced with the lentiCas9-Blast (Addgene #52962) and successfully transduced cells were selected for using culture medium supplemented with 10 µg/mL blasticidin. Single cell clones were generated, and the Cas9 editing efficiency was assessed using flow cytometry following transduction with lentivirus generated using lentivirus generated with pKLV2-U6-gRNA5(gBFP)-PGKGFP2ABFP-W (Addgene #67984) (Supplementary Figure 1).

To perform genome-wide CRISPR/Cas9 screens, 1.2 × 10^8^ CLEC213-Cas9 cells were transduced with lentivirus containing the chicken sgRNA library at a multiplicity of infection (MOI) of 0.3 to ensure majority of cells receive no more than one sgRNA per cell according to Poisson distribution, with at least 500 cells for each sgRNA. The cells were selected with 2 µg/mL puromycin (Invivogen) for 10 days to achieve >95% gene disruption. 4.0 × 10^7^ CLEC213-GeCKO library cells were subjected to deep sequencing to confirm the library integrity.

3 × 10^8^ CLEC213-GeCKO library cells were infected with MOI 3 for UDL or MOI 5 for PR8 to achieve >95% infected cells. The cells were harvested and fixed in 4% paraformaldehyde (ThermoScientific) 12 hpi. Cells were stained with anti-Influenza A M2 (14c2) AF488 antibody (sc-32238-AF488) and analysed using the BD FACs Aria flow cytometer to separate the cells into a low, mid and high population based on M2 staining. The “low” and “high” populations contained the bottom and top 5% M2 stained cells respectively, and the “mid” population contained the middle 10% stained cells.

The genomic DNA (gDNA) from each FACS sorted population (low, mid and high) was extracted using QIAamp DNA FFPE Tissue kit (Qiagen) according to the manufacturer’s instructions. The inserted gRNA cassette was amplified using a Q5 high-fidelity Hot start DNA polymerase (NEB) using primers outlined in Supplementary Table 1. PCR fragments were analysed using gel electrophoresis before the PCR products were purified using PCR purification kits (Qiagen). A second PCR reaction using NEBNext Ultra II Q5 polymerase (NEB) was performed to incorporate sequencing bar codes (Supplementary Table 1) and the PCR products were purified using AMPure XP beads and sequenced using Illumina NextSeq2000 platform (Edinburgh Clinical Research Facility).

Raw data were processed to eliminate reads of low quality or incorrect guide sequence. The reads matching the sgRNA sequence were selected with a python script. Using the MAGeCK analysis pipeline [35], the enrichment of genes in either the “low” or “high” sorted populations was compared to the “mid” population. Candidate hits were assessed based on the enriched reads number for each guide; how many guides per gene were similarly enriched; and to what degree they were enriched in the UDL and PR8 screens.

Gene ontology (GO) analysis was performed using ShinyGO 0.77 software. The gene list from the chicken library was uploaded to the background as recommended. Enriched biological processes (FDR<0.05) in the pathway database were analysed using HDFs with *P*<0.01 as the selection criteria.

### Generation of CRISPR KO cell lines and single cell clones

Non-targeting control gRNA and gRNAs targeting SLC35A1, ANP32A, MOGS, MGAT1, DENR, DMXL1, PTAR1, IPO9, KLF6, ENO1 and TSG101 were cloned into the BbsI-digested pKLV2-U6-gRNA5(BbsI)-PGKpuro2ABFP-W plasmid individually. Lentivirus was generated by transfection of HEK293T cells with guide inserted into pKLV2-U6-gRNA5(BbsI)-PGKpuro2ABFP-W plasmid and the packaging plasmids psAX2 and pMD2.G with lipofectamine 2000 (ThermoFisher) as per the manufacturer’s protocol. Lentivirus was harvested after 72 hours incubation at 37°C 5% CO_2_. To generate polyclonal CRISPR KO cells, the generated lentivirus was transduced into the appropriate Cas9-expressing cell line with 0.16 µg/mL dextran (Supplementary Table 2). Mutagenesis was confirmed by gDNA isolation using the QIAamp DNA FFPE Tissue kit followed by PCR amplification of the edited region, subsequent PCR product purification, Sanger Sequencing and TIDE analysis where possible (Supplementary Table 3).

### Virus infection and replication in different cells

To determine the effect of KO on viral infection in selected KO cell lines by flow cytometry, cells were infected with a high MOI of virus as indicated for 45 minutes in serum-free medium (SFM). SFM supplemented with 0.14% bovine serum albumin (BSA) fraction V was added to each well and the cells collected and fixed with 2% paraformaldehyde 16 hpi.

To determine the effect on viral infection in selected KO cell lines by virus titration, cells were infected at the specified MOI with either PR8 or UDL. Cells were washed 1 hpi and medium replaced with viral growth medium (VGM; SFM supplemented with 0.14% BSA fraction V and 0.2 µg/mL TPCK) for incubation at 37°C 5% CO_2_. VGM for DF-1 cells contained 0.5 µg/mL TPCK, and VGM for A549 and MDCK cells contained 1 µg/mL TPCK.

Supernatants for UDL and PR8 infections were collected at the times indicated in the text, serially diluted and titred on MDCK cells using VGM and 1.2% cellulose at its final concentration for 48 hours. Supernatants for H5N2 (A/chicken/Belgium/150/1999), H6N1 (A/chicken/Netherlands/917/2010) and H9N2 (A/chicken/Saudi Arabia/2525/2000) infections were collected at the times indicated in the text, serially diluted and titred on MDCK_chANP32A^+^ cells using VGM and 0.8% cellulose at its final concentration for 48 hours.

### Immunofluorescence staining

To determine the percentage of cells infected in each population, infected cells were harvested and fixed with 4% paraformaldehyde. For H1Nx infected cells, samples were blocked in 3% FBS/PBS (v/v) prior to staining with 50 µg/mL Anti-M2-488 (SantaCruz). For H5N2, H6N1 and H9N2 infected cells, samples were permeabilised in 0.1% Triton X-100 prior to blocking with 3% FBS/PBS (v/v). Samples were then stained with 2 µg/mL mouse anti-NP monoclonal antibody AA5H (Invitrogen) which was labelled using the Alexa Fluor 647 Conjugation Kit – Lightning Link (AbCam). Samples were analysed using the BD Fortessa flow cytometer.

To detect SA expression on the surface of cells, cells were cultured on coverslips and fixed in 2% paraformaldehyde and washed in PBS containing 0.05% Tween20 (v/v) (PBS-T). Samples were stained with 2 µg/mL biotinylated MAL II (Maackia amurensis lectin; Vector laboratories) for α2,3-linked SA expression or biotinylated SNA (Sambucus nigra lectin; Vector Laboratories) for α2,6-linked SA expression. Samples were then washed with PBS-T and stained in 1 µg/mL streptavidin-488 (AAT Bioquest) before being washed with PBS-T and counterstained with DAPI before mounting coverslips on microscope slides with ProLong gold (Invitrogen). Samples were imaged using the LSM 880 confocal microscope. Samples stained with anti-biotinylated SNA antibody were altered from the green channel to the red channel post-imaging for visual clarity between MAL II^+^ (green) and SNA^+^ (red) cells.

### Quantification and statistical analysis

Quantitative data are presented as means ± standard deviations (SDs) of three independent experiments or replicates. Data were statistically analysed using the appropriate statistical test as specified for each experiment using GraphPad Prism software.

## Acknowledgements

We would like to thank Dr. Colin Sharp (The Roslin Institute, University of Edinburgh) for the mouse anti-NP-647 conjugate monoclonal antibody, Dr. Sascha Trapp (French National Institute for Agriculture, Food and Environment, France) for CLEC213 cells, Prof. John Doench for the advice and expertise offered over the course of this work, Prof. Massimo Palmarini (MRC-University of Glasgow Centre for Virus Research, UK) for MDCK/chicken ANP32A cells and Prof. Sjaak de Wit (R&D department of Royal GD, Deventer, Utrecht University) for avian IAV strains.

We are grateful to the staff of the Roslin Institute’s Central Services Unit, Facilities team and Bioimaging facilities.

This work was funded by Cobb-Vantress, BBSRC BBS/E/RL/230002A from the Biotechnology and Biological Sciences Research Council (BBSCRC) to FG, PD and SE2223 from the BBSRC under the aegis of the International Coordination of Research on Infectious Animal Diseases (ICRAD, SE2223) programme to FG and PD.

## Supplementary Materials

**Supplementary Figure 1.**
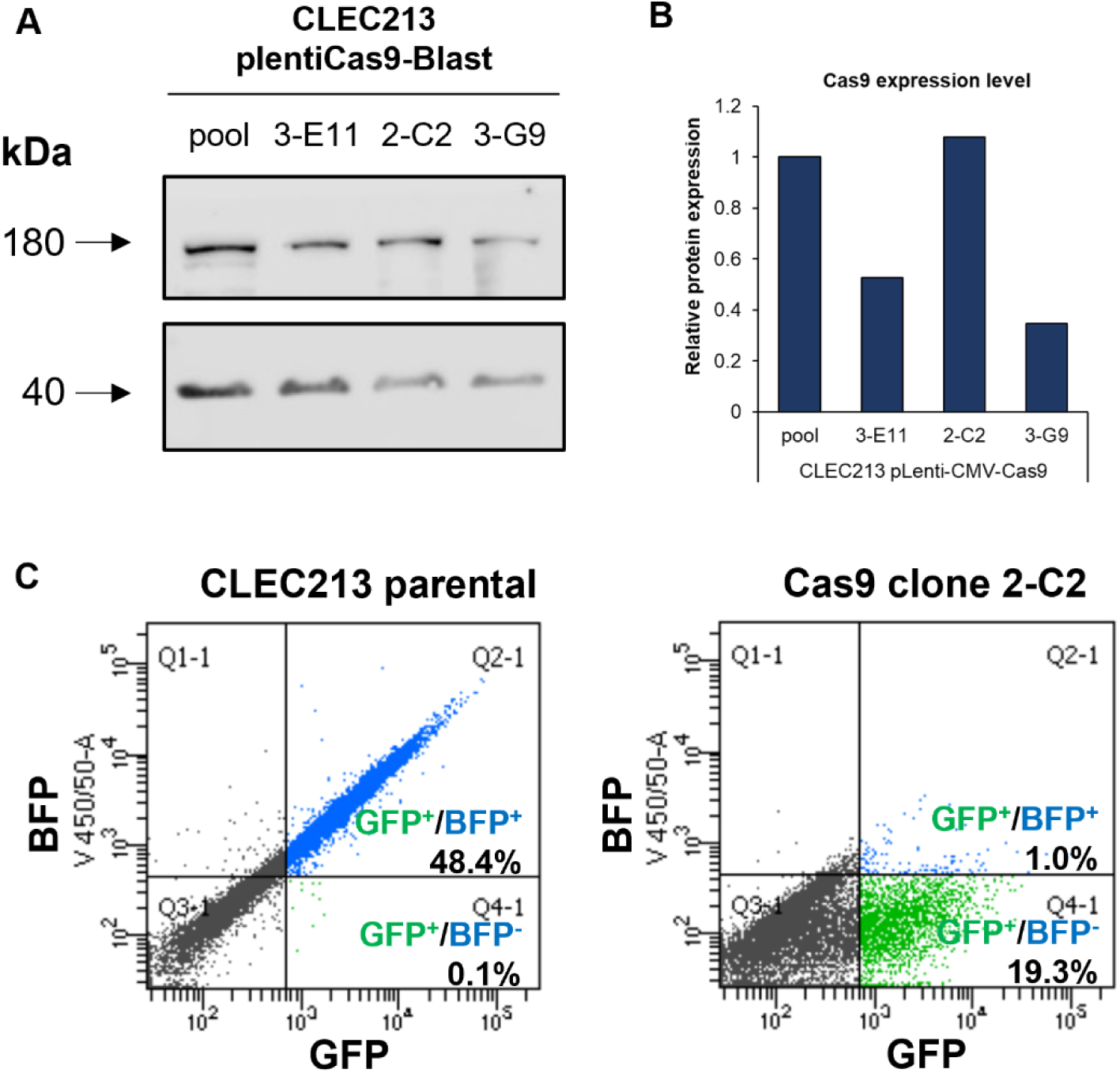
Selection and characterisation of CLEC213-Cas9 single cell clone. (A) Western blot detecting expression of Cas9 protein (180 kDa) and actin (40 kDa) as a loading control for three single cell clones. (B) Quantification of Cas9 expression normalised to actin control. (c) Flow cytometry of CLEC213 parental and selected 2-C2 CLEC213-Cas9 single transduced with pKLV2 lentivirus encoding BFP/GFP reporters and a BFP sequence in the sgRNA site.

**Supplementary Figure 2.**
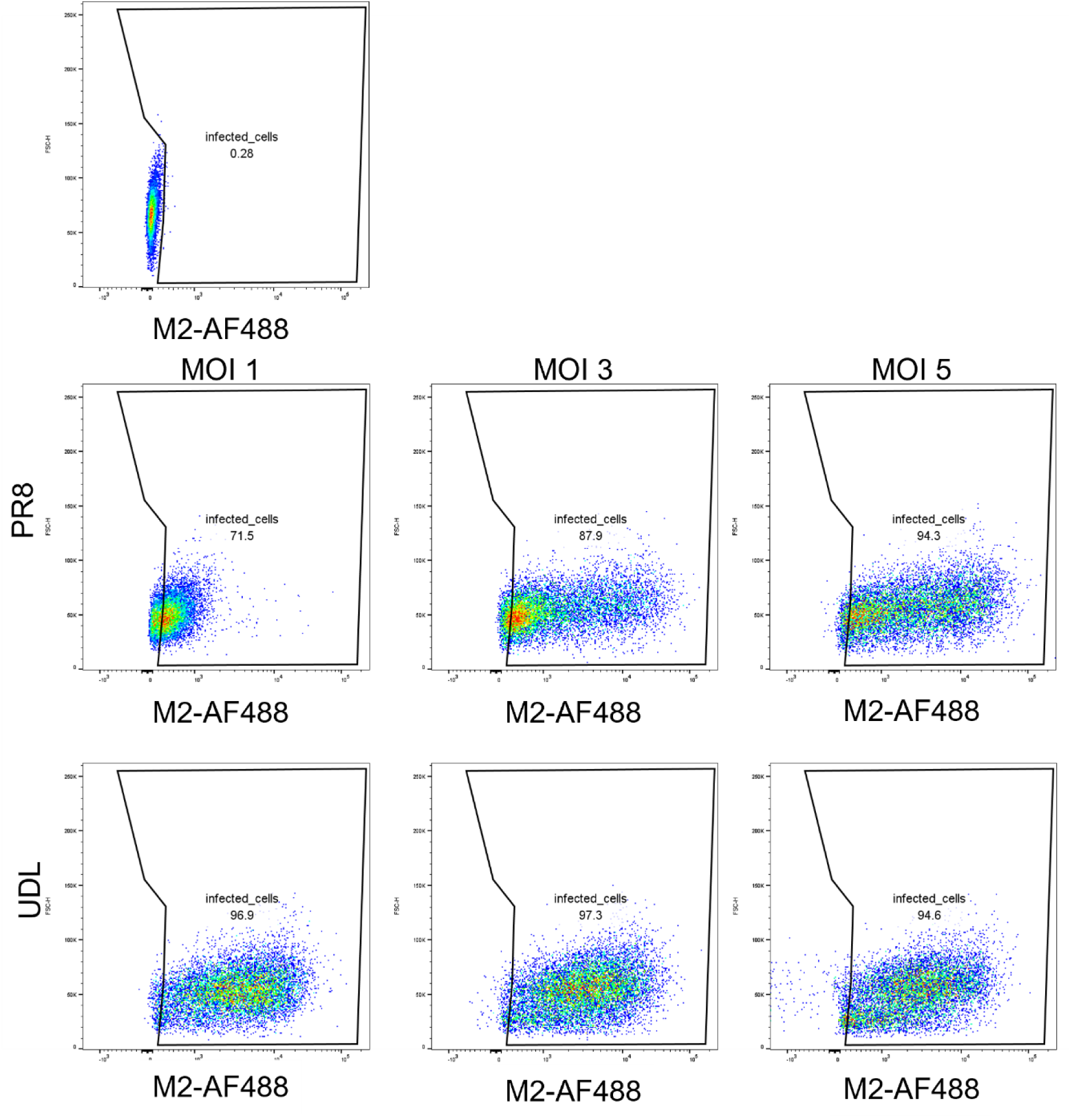
MOI test to establish infection of ∼95% of CLEC213 cells by flow cytometry. Cells were infected with PR8 or UDL 3:5 reassortant virus at MOI 1, 3 or 5 for 16 hours. Flow cytometry was used to determine the number of infected cells post anti-M2-AF488 antibody staining.

**Supplementary Figure 3.**
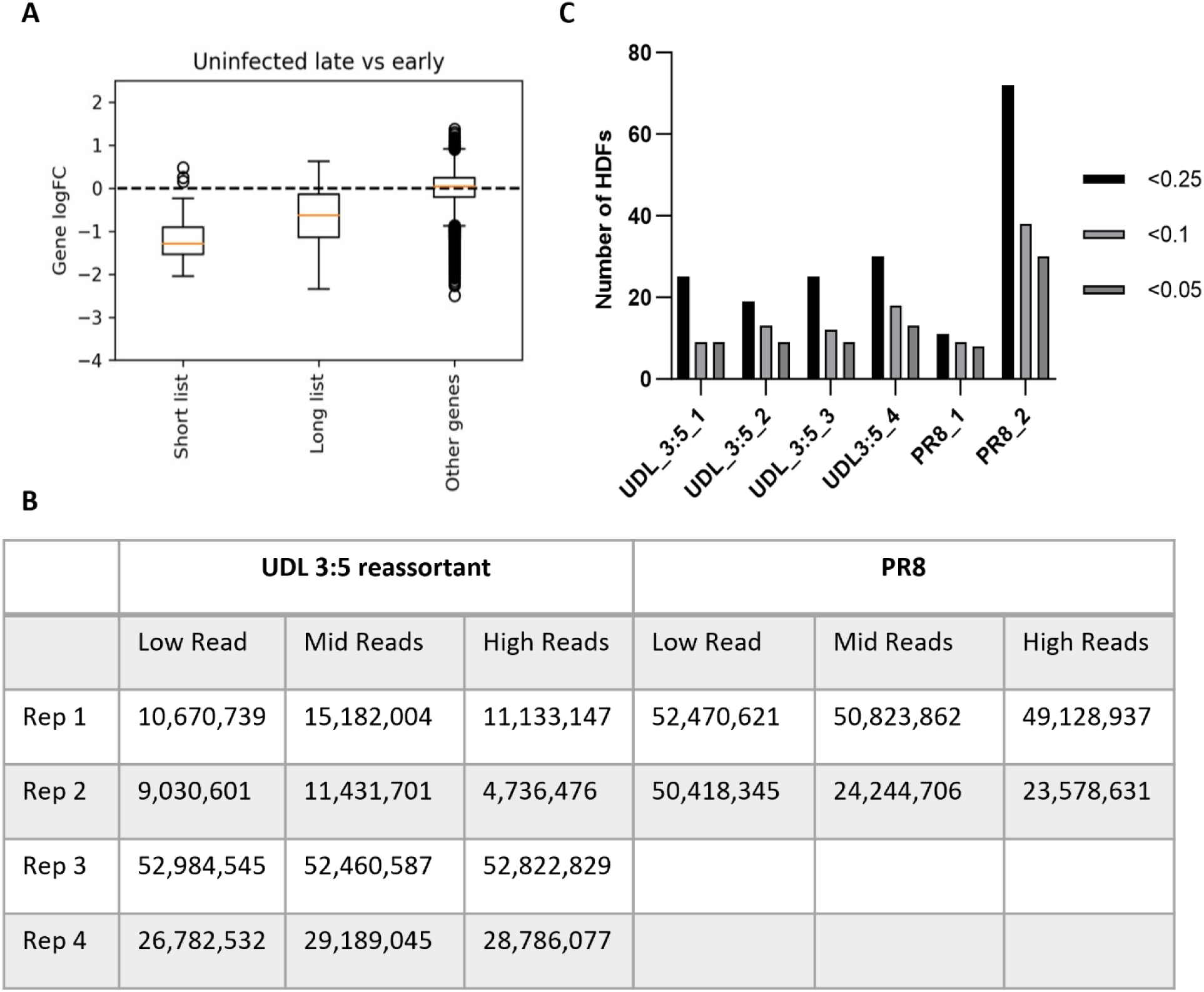
Analysis of sgRNA representation in the CLEC213-GeCKO library and post IAV infection. (A) Box plot displaying the drop-out of essential genes in uninfected transduced cells. (B) Summary of Illumina reads in each M2 stained sorted population after FastQC analysis. (C) Number of HDFs identified at various FDR thresholds for each screen.

**Supplementary Figure 4.**
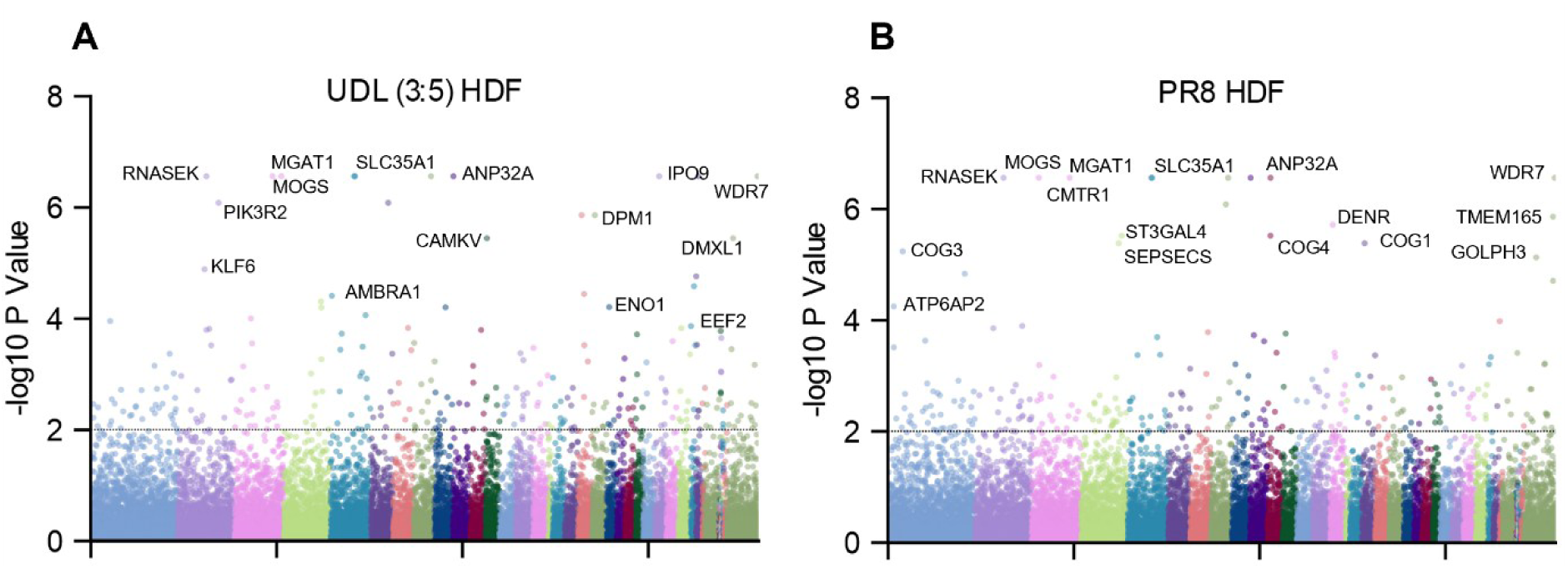
Spatial distribution of CRISPR KO HDFs on the genome. Host dependency identified in the (A) UDL 3:5 reassortant screen and the (B) PR8 screen. Gene-level –log10 *P* values are plotted (*y-*axis) against the chromosome the genes are located in (*x*-axis). A selection of top hits are highlighted.

**Supplementary Figure 5.**
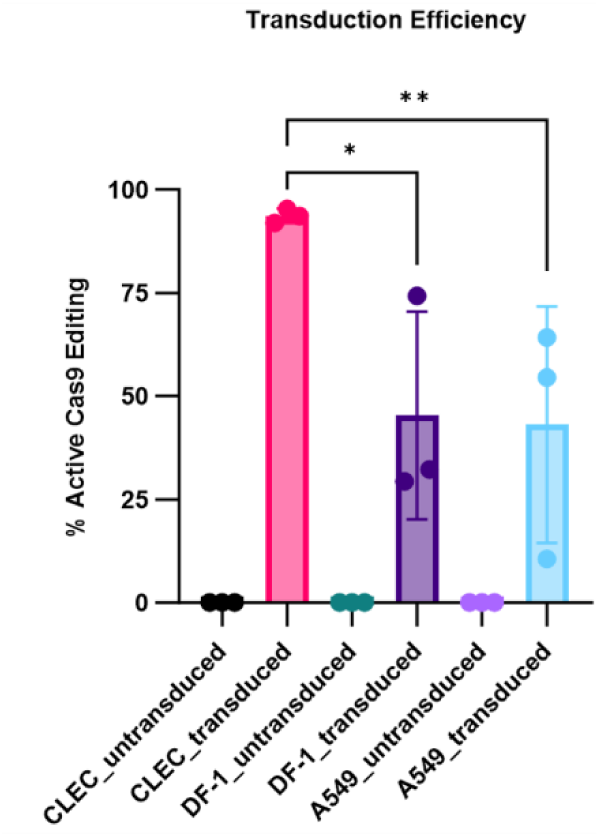
Comparison of the Cas9 editing efficiency between different cell lines. CLEC213, DF-1 and A549 cells expressing Cas9 were transduced with a lentivirus reporter (pKLV2_U6(BFP)BFP_GFP). Post transduction, cells were fixed and BFP:GFP was measured using a flow cytometer. Percentage of Cas9 editing cells was calculated by BFP-GFP+/total BFP-GFP+ and BFP+GFP+ cells*100.

**Supplementary Figure 6.**
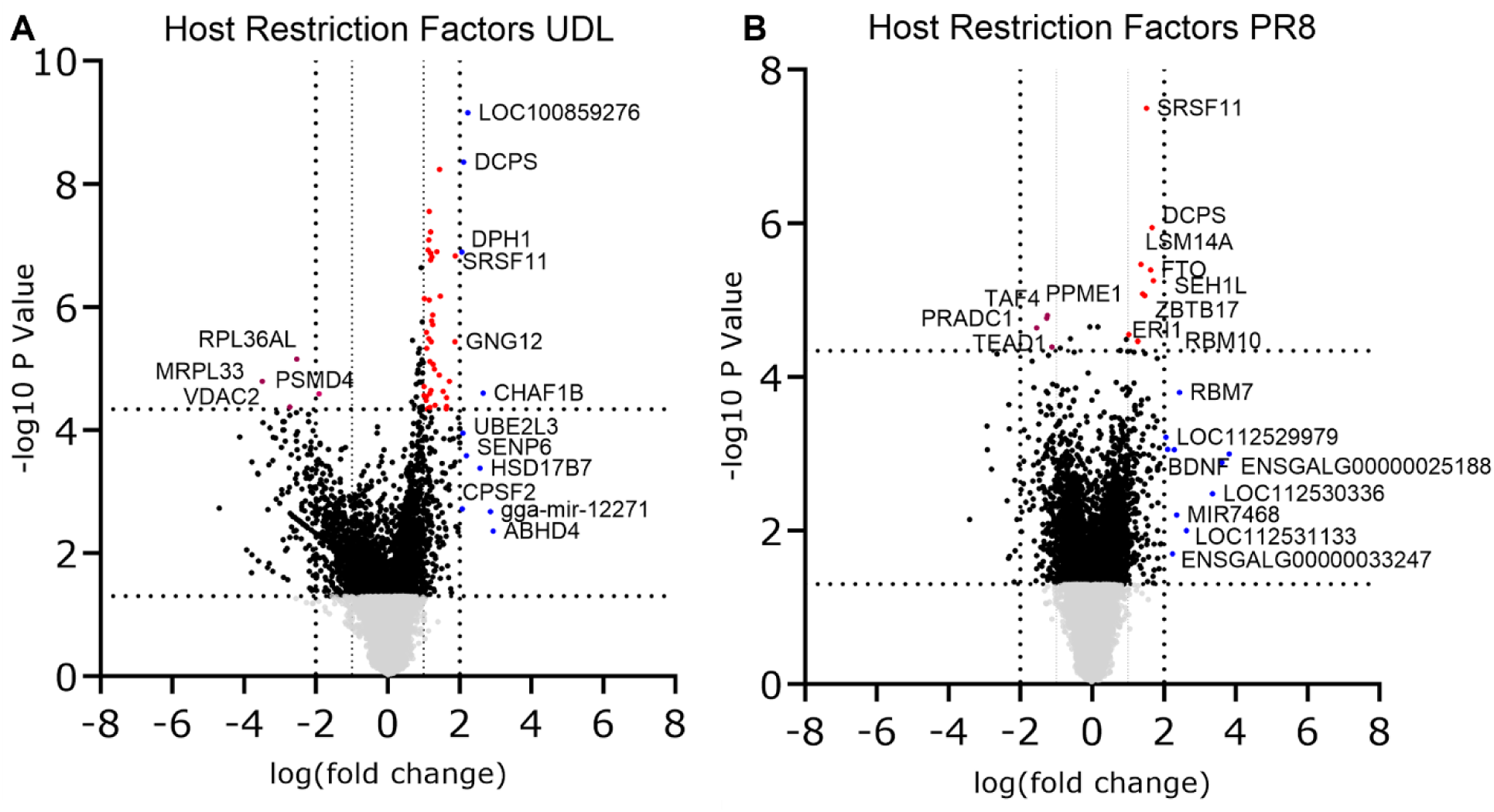
Host restriction factors identified in the UDL and PR8 sort screens. MAGeCK analysis of multiple replicates comparing guides enriched in the “high” M2+ population to the “mid” M2+ population yielded log fold changes that were plotted on the *x*-axis and negative log10 P values that were plotted on the *y*-axis for both UDL 3:5 reassortant (A) and PR8 (B) screens. A selection of hits have been labelled.

**Supplementary Table 1.**
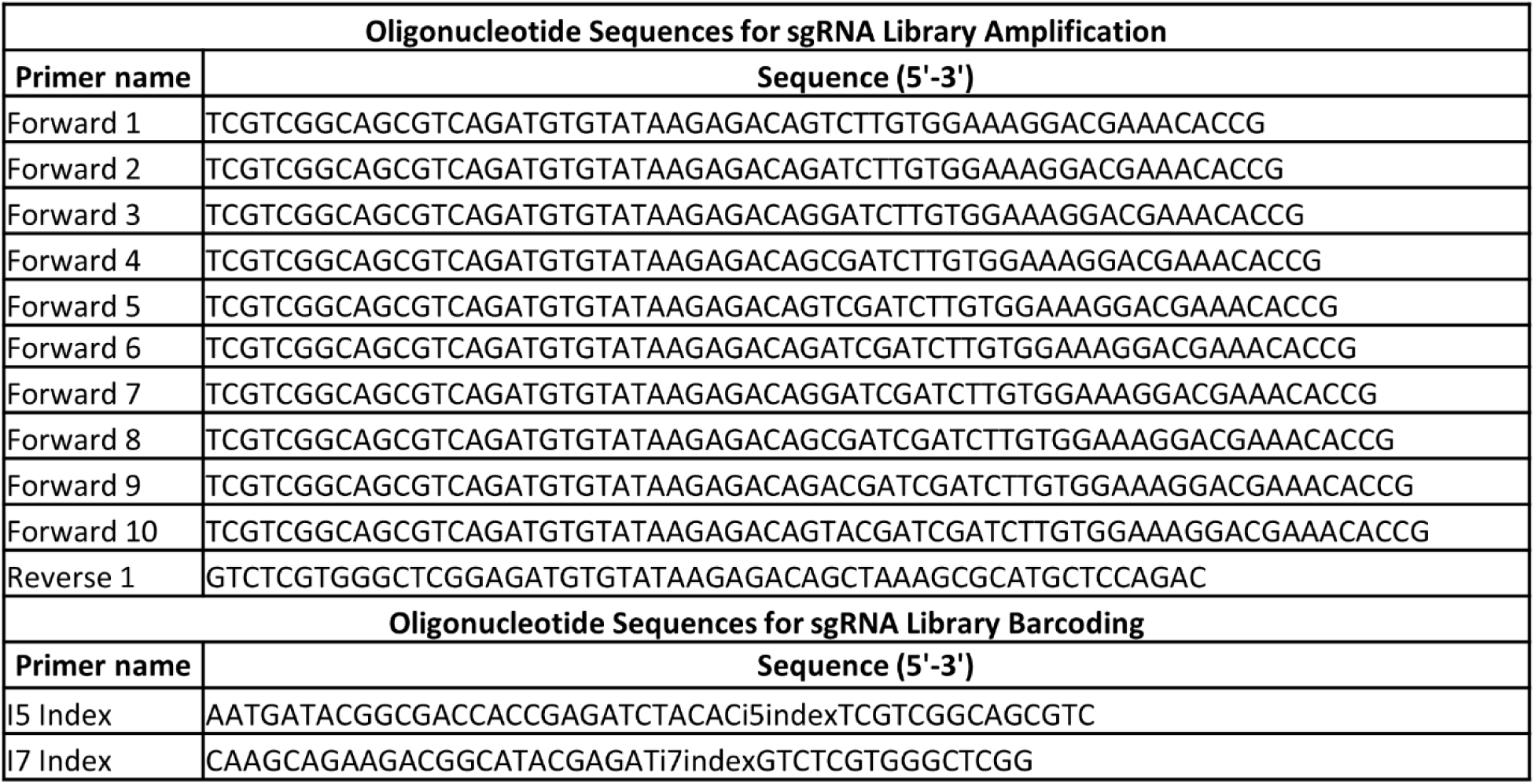
Oligonucleotide sequences for amplifying and sequencing the sgRNA library from cells.

**Supplementary Table 2.**
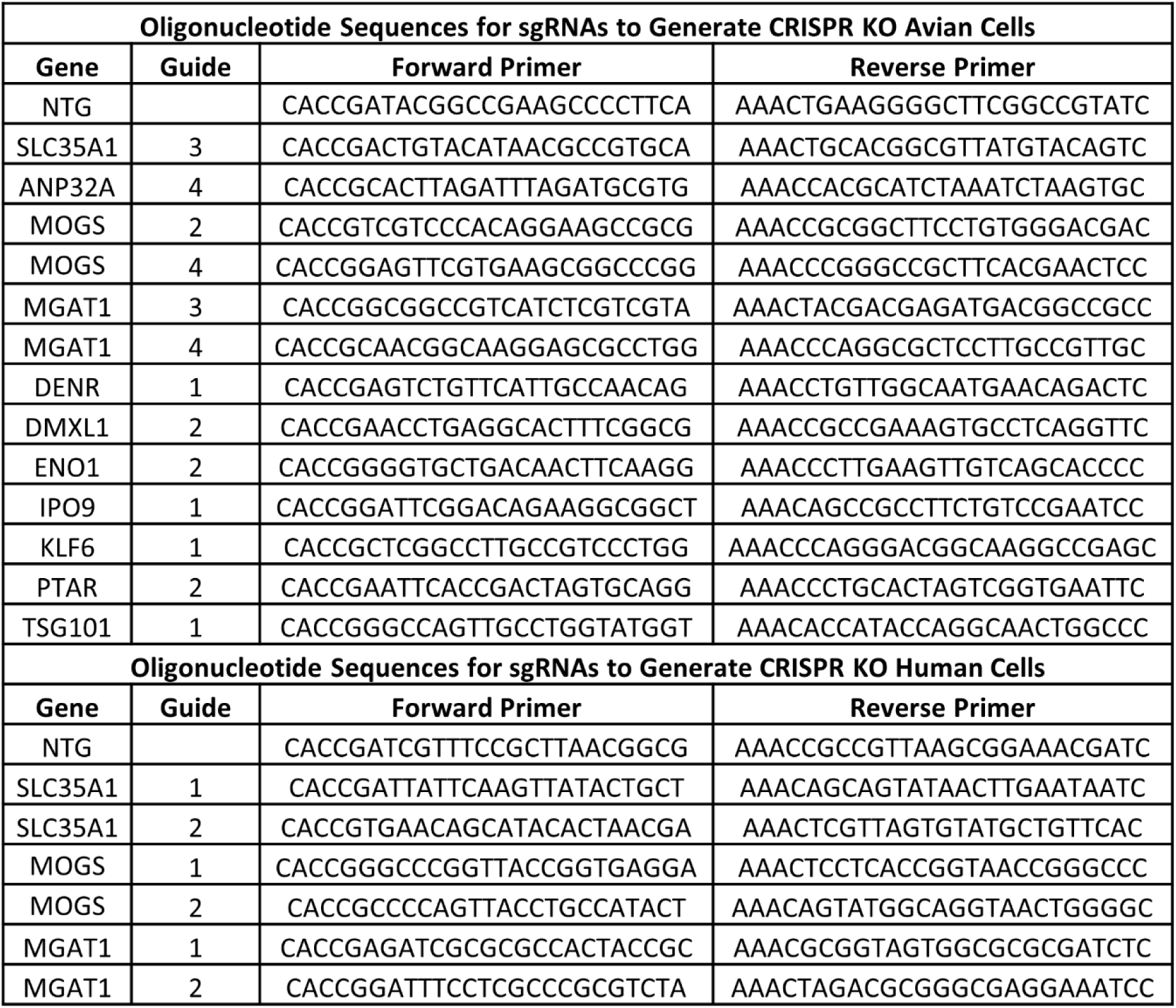
Oligonucleotide sequences to generate KOs in selected genes in avian (CLEC213 and DF-1) or human (A549) cell lines.

**Supplementary Table 3.**
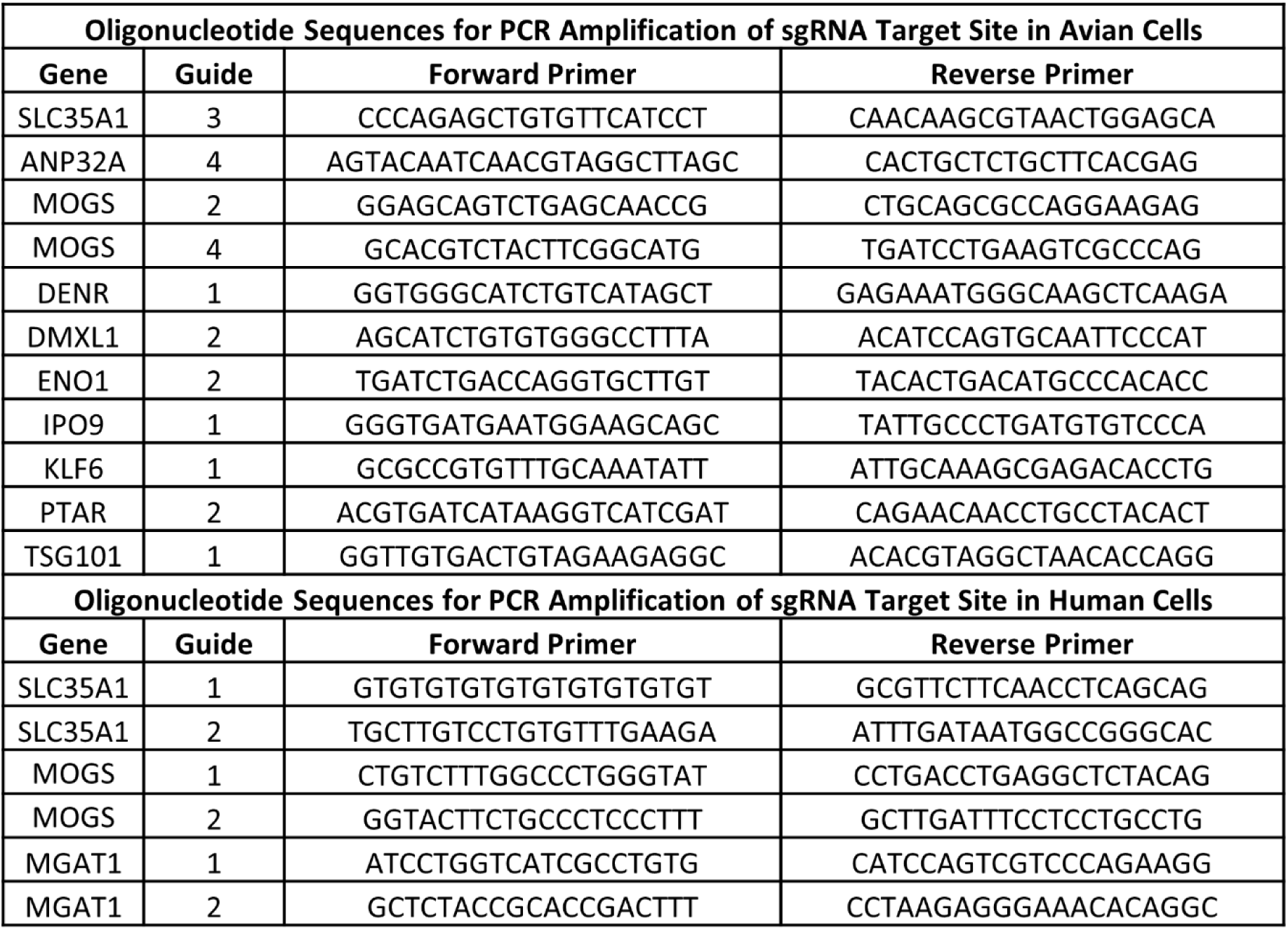
Oligonucleotide sequences for amplifying the sgRNA target site in in avian (CLEC213 and DF-1) or human (A549) cell lines to confirm KO of targeted gene.

