## Supplementary Figures for "Identification of key host genes for influenza A virus in avian cells using a genome-wide CRISPR-Cas9 screen"

Science Advances

AAAS

For papers with only two authors: Rosemary Anna Blake *et al.*

**This PDF file includes:**

Figs. S1 to S6
Tables S1 to S3

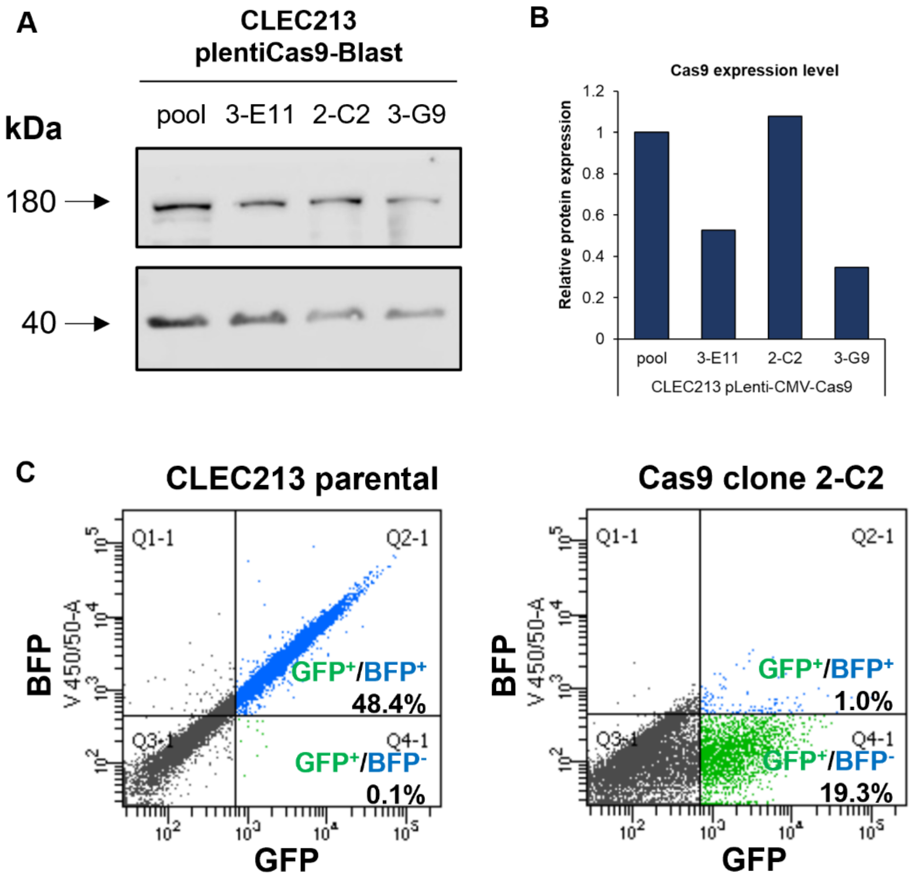


**Supplementary Figure 1| Selection and characterisation of CLEC213-Cas9 single cell clone.** (A) Western blot detecting expression of Cas9 protein (180 kDa) and actin (40 kDa) as a loading control for three single cell clones. (B) Quantification of Cas9 expression normalised to actin control. (c) Flow cytometry of CLEC213 parental and selected 2-C2 CLEC213-Cas9 single transduced with pKLV2 lentivirus encoding BFP/GFP reporters and a BFP sequence in the sgRNA site.

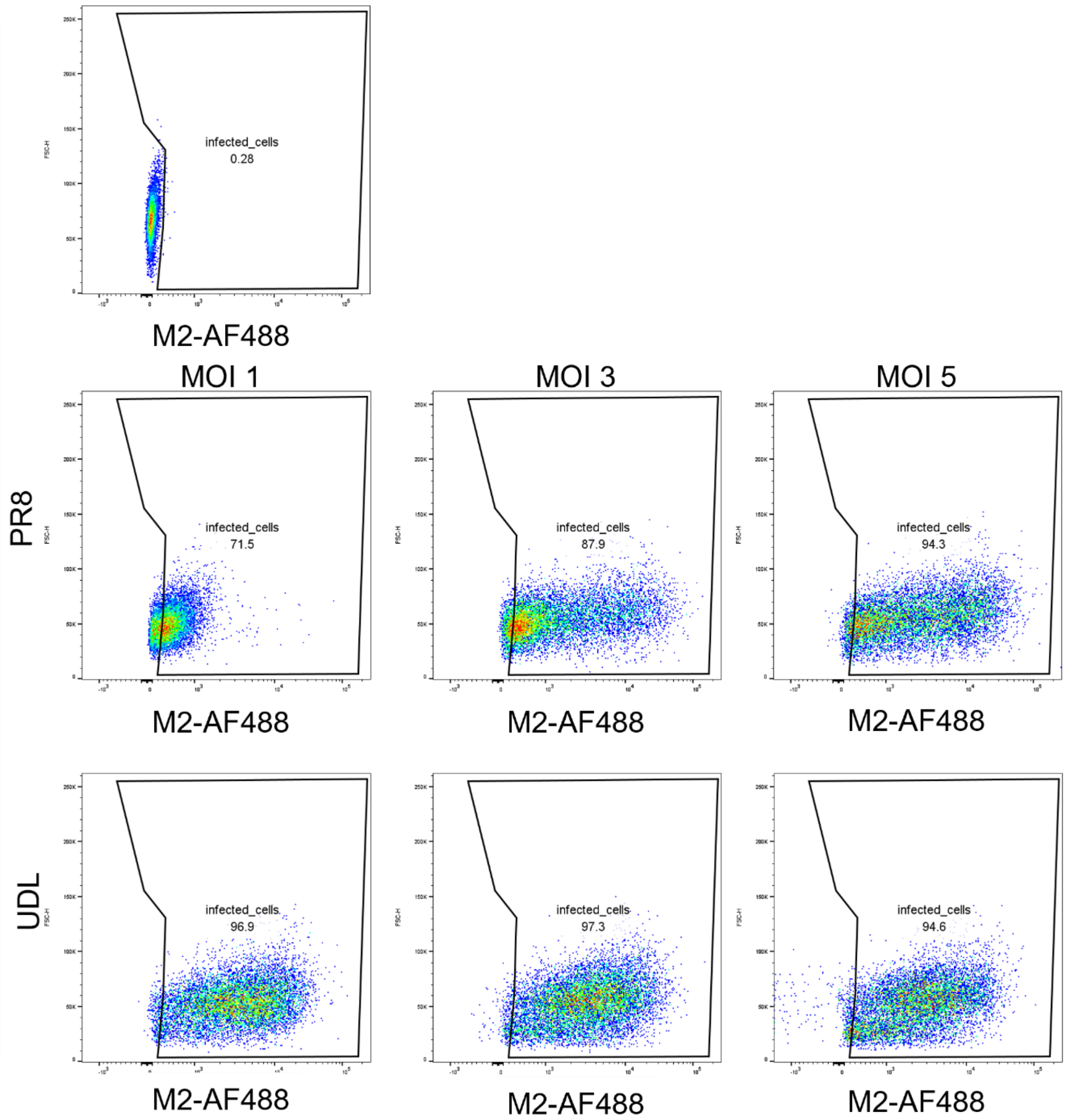


**Supplementary Figure 2| MOI test to establish infection of ~95% of CLEC213 cells by flow cytometry.** Cells were infected with PR8 or UDL 3:5 reassortant virus at MOI 1, 3 or 5 for 16 hours. Flow cytometry was used to determine the number of infected cells post anti-M2-AF488 antibody staining.

A C

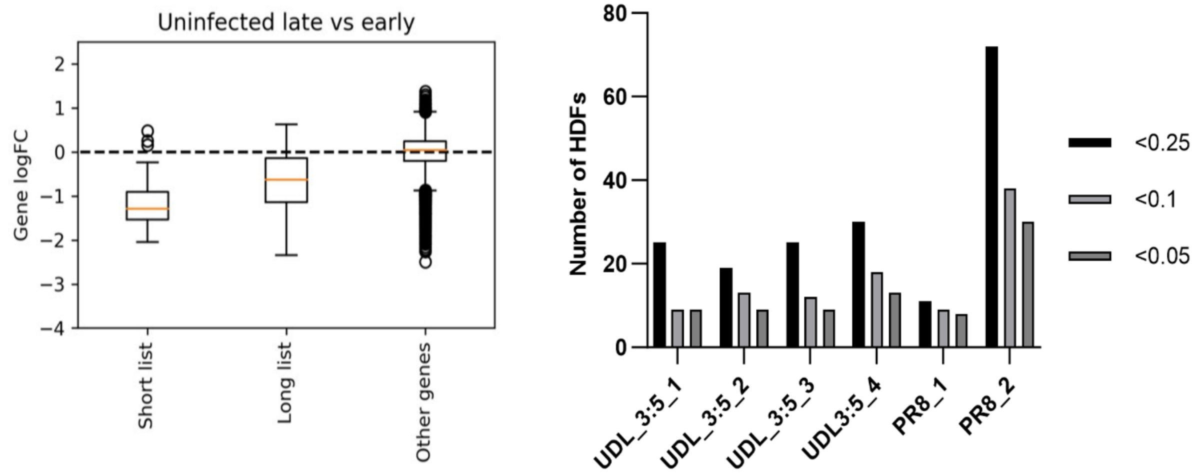


B

| | UDL 3:5 reassortant | | | PR8 | | |
|---|---|---|---|---|---|---|
| | Low Read | Mid Reads | High Reads | Low Read | Mid Reads | High Reads |
| Rep 1 | 10,670,739 | 15,182,004 | 11,133,147 | 52,470,621 | 50,823,862 | 49,128,937 |
| Rep 2 | 9,030,601 | 11,431,701 | 4,736,476 | 50,418,345 | 24,244,706 | 23,578,631 |
| Rep 3 | 52,984,545 | 52,460,587 | 52,822,829 | | | |
| Rep 4 | 26,782,532 | 29,189,045 | 28,786,077 | | | |

**Supplementary Figure 3| Analysis of sgRNA representation in the CLEC213-GeCKO library and post IAV infection.** (A) Box plot displaying the drop-out of essential genes in uninfected transduced cells. (B) Summary of Illumina reads in each M2 stained sorted population after FastQC analysis. (C) Number of HDFs identified at various FDR thresholds for each screen.

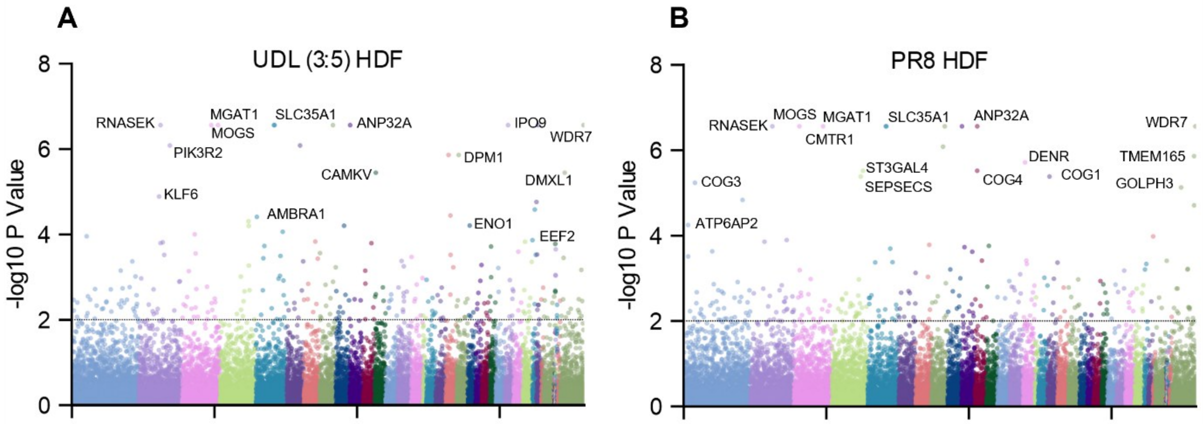


**Supplementary Figure 4| Spatial distribution of CRISPR KO HDFs on the genome.** Host dependency identified in the (A) UDL 3:5 reassortant screen and the (B) PR8 screen. Gene-level –log10 *P* values are plotted (*y*-axis) against the chromosome the genes are located in (*x*-axis). A selection of top hits are highlighted.

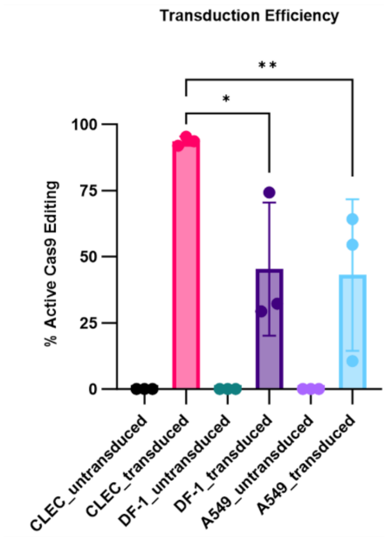


**Supplementary Figure 5| Comparison of the Cas9 editing efficiency between different cell lines.** CLEC213, DF-1 and A549 cells expressing Cas9 were transduced with a lentivirus reporter (pKLV2_U6(BFP)BFP_GFP). Post transduction, cells were fixed and BFP:GFP was measured using a flow cytometer. Percentage of Cas9 editing cells was calculated by BFP-GFP+/total BFP-GFP+ and BFP+GFP+ cells*100.

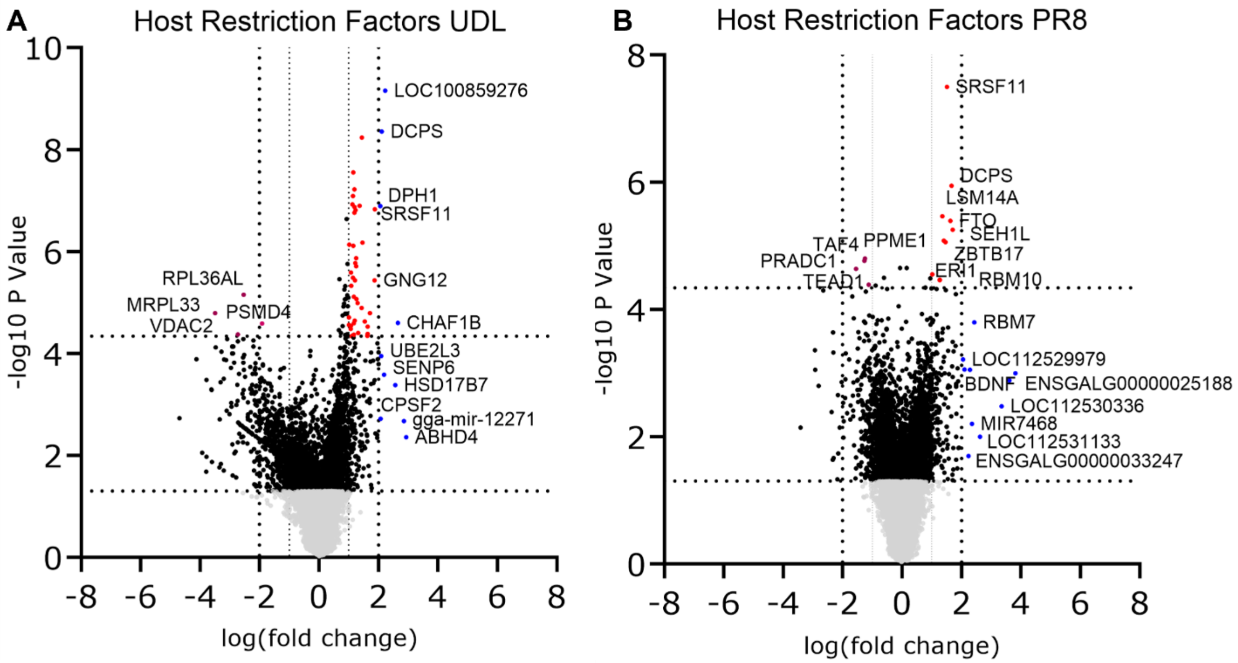


**Supplementary Figure 6| Host restriction factors identified in the UDL and PR8 sort screens.** MAGeCK analysis of multiple replicates comparing guides enriched in the "high" M2+ population to the "mid" M2+ population yielded log fold changes that were plotted on the *x*-axis and negative log10 P values that were plotted on the *y*-axis for both UDL 3:5 reassortant (A) and PR8 (B) screens. A selection of hits have been labelled.

| Oligonucleotide Sequences for sgRNA Library Amplification | |
|---|---|
| **Primer name** | **Sequence (5'-3')** |
| Forward 1 | TCGTCGGCAGCGTCAGATGTGTATAAGAGACAGTCTTGTGGAAAGGACGAAACACCG |
| Forward 2 | TCGTCGGCAGCGTCAGATGTGTATAAGAGACAGATCTTGTGGAAAGGACGAAACACCG |
| Forward 3 | TCGTCGGCAGCGTCAGATGTGTATAAGAGACAGGATCTTGTGGAAAGGACGAAACACCG |
| Forward 4 | TCGTCGGCAGCGTCAGATGTGTATAAGAGACAGCGATCTTGTGGAAAGGACGAAACACCG |
| Forward 5 | TCGTCGGCAGCGTCAGATGTGTATAAGAGACAGTCGATCTTGTGGAAAGGACGAAACACCG |
| Forward 6 | TCGTCGGCAGCGTCAGATGTGTATAAGAGACAGATCGATCTTGTGGAAAGGACGAAACACCG |
| Forward 7 | TCGTCGGCAGCGTCAGATGTGTATAAGAGACAGGATCGATCTTGTGGAAAGGACGAAACACCG |
| Forward 8 | TCGTCGGCAGCGTCAGATGTGTATAAGAGACAGCGATCGATCTTGTGGAAAGGACGAAACACCG |
| Forward 9 | TCGTCGGCAGCGTCAGATGTGTATAAGAGACAGACGATCGATCTTGTGGAAAGGACGAAACACCG |
| Forward 10 | TCGTCGGCAGCGTCAGATGTGTATAAGAGACAGTACGATCGATCTTGTGGAAAGGACGAAACACCG |
| Reverse 1 | GTCTCGTGGGCTCGGAGATGTGTATAAGAGACAGCTAAAGCGCATGCTCCAGAC |
| **Oligonucleotide Sequences for sgRNA Library Barcoding** | |
| **Primer name** | **Sequence (5'-3')** |
| I5 Index | AATGATACGGCGACCACCGAGATCTACACi5indexTCGTCGGCAGCGTC |
| I7 Index | CAAGCAGAAGACGGCATACGAGATi7indexGTCTCGTGGGCTCGG |

**Supplementary Table 1| Oligonucleotide sequences for amplifying and sequencing the sgRNA library from cells.**

| Oligonucleotide Sequences for sgRNAs to Generate CRISPR KO Avian Cells | | | |
|---|---|---|---|
| **Gene** | **Guide** | **Forward Primer** | **Reverse Primer** |
| NTG | | CACCGATACGGCCGAAGCCCCTTCA | AAACTGAAGGGGCTTCGGCCGTATC |
| SLC35A1 | 3 | CACCGACTGTACATAACGCCGTGCA | AAACTGCACGGCGTTATGTACAGTC |
| ANP32A | 4 | CACCGCACTTAGATTTAGATGCGTG | AAACCACGCATCTAAATCTAAGTGC |
| MOGS | 2 | CACCGTCGTCCCACAGGAAGCCGCG | AAACCGCGGCTTCCTGTGGGACGAC |
| MOGS | 4 | CACCGGAGTTCGTGAAGCGGCCCGG | AAACCCGGGCCGCTTCACGAACTCC |
| MGAT1 | 3 | CACCGGCGGCCGTCATCTCGTCGTA | AAACTACGACGAGATGACGGCCGCC |
| MGAT1 | 4 | CACCGCAACGGCAAGGAGCGCCTGG | AAACCCAGGCGCTCCTTGCCGTTGC |
| DENR | 1 | CACCGAGTCTGTTCATTGCCAACAG | AAACCTGTTGGCAATGAACAGACTC |
| DMXL1 | 2 | CACCGAACCTGAGGCACTTTCGGCG | AAACCGCCGAAAGTGCCTCAGGTTC |
| ENO1 | 2 | CACCGGGGTGCTGACAACTTCAAGG | AAACCCTTGAAGTTGTCAGCACCCC |
| IPO9 | 1 | CACCGGATTCGGACAGAAGGCGGCT | AAACAGCCGCCTTCTGTCCGAATCC |
| KLF6 | 1 | CACCGCTCGGCCTTGCCGTCCCTGG | AAACCCAGGGACGGCAAGGCCGAGC |
| PTAR | 2 | CACCGAATTCACCGACTAGTGCAGG | AAACCCTGCACTAGTCGGTGAATTC |
| TSG101 | 1 | CACCGGGCCAGTTGCCTGGTATGGT | AAACACCATACCAGGCAACTGGCCC |

| Oligonucleotide Sequences for sgRNAs to Generate CRISPR KO Human Cells | | | |
|---|---|---|---|
| **Gene** | **Guide** | **Forward Primer** | **Reverse Primer** |
| NTG | | CACCGATCGTTTCCGCTTAACGGCG | AAACCGCCGTTAAGCGGAAACGATC |
| SLC35A1 | 1 | CACCGATTATTCAAGTTATACTGCT | AAACAGCAGTATAACTTGAATAATC |
| SLC35A1 | 2 | CACCGTGAACAGCATACACTAACGA | AAACTCGTTAGTGTATGCTGTTCAC |
| MOGS | 1 | CACCGGGCCCGGTTACCGGTGAGGA | AAACTCCTCACCGGTAACCGGGCCC |
| MOGS | 2 | CACCGCCCCAGTTACCTGCCATACT | AAACAGTATGGCAGGTAACTGGGGC |
| MGAT1 | 1 | CACCGAGATCGCGCGCCACTACCGC | AAACGCGGTAGTGGCGCGCGATCTC |
| MGAT1 | 2 | CACCGGATTTCCTCGCCCGCGTCTA | AAACTAGACGCGGGCGAGGAAATCC |

**Supplementary Table 2| Oligonucleotide sequences to generate KOs in selected genes in avian (CLEC213 and DF-1) or human (A549) cell lines.**

| Oligonucleotide Sequences for PCR Amplification of sgRNA Target Site in Avian Cells | | | |
|---|---|---|---|
| **Gene** | **Guide** | **Forward Primer** | **Reverse Primer** |
| SLC35A1 | 3 | CCCAGAGCTGTGTTCATCCT | CAACAAGCGTAACTGGAGCA |
| ANP32A | 4 | AGTACAATCAACGTAGGCTTAGC | CACTGCTCTGCTTCACGAG |
| MOGS | 2 | GGAGCAGTCTGAGCAACCG | CTGCAGCGCCAGGAAGAG |
| MOGS | 4 | GCACGTCTACTTCGGCATG | TGATCCTGAAGTCGCCCAG |
| DENR | 1 | GGTGGGCATCTGTCATAGCT | GAGAAATGGGCAAGCTCAAGA |
| DMXL1 | 2 | AGCATCTGTGTGGGCCTTTA | ACATCCAGTGCAATTCCCAT |
| ENO1 | 2 | TGATCTGACCAGGTGCTTGT | TACACTGACATGCCCACACC |
| IPO9 | 1 | GGGTGATGAATGGAAGCAGC | TATTGCCCTGATGTGTCCCA |
| KLF6 | 1 | GCGCCGTGTTTGCAAATATT | ATTGCAAAGCGAGACACCTG |
| PTAR | 2 | ACGTGATCATAAGGTCATCGAT | CAGAACAACCTGCCTACACT |
| TSG101 | 1 | GGTTGTGACTGTAGAAGAGGC | ACACGTAGGCTAACACCAGG |
| **Oligonucleotide Sequences for PCR Amplification of sgRNA Target Site in Human Cells** | | | |
| **Gene** | **Guide** | **Forward Primer** | **Reverse Primer** |
| SLC35A1 | 1 | GTGTGTGTGTGTGTGTGTGT | GCGTTCTTCAACCTCAGCAG |
| SLC35A1 | 2 | TGCTTGTCCTGTGTTTGAAGA | ATTTGATAATGGCCGGGCAC |
| MOGS | 1 | CTGTCTTTGGCCCTGGGTAT | CCTGACCTGAGGCTCTACAG |
| MOGS | 2 | GGTACTTCTGCCCTCCCTTT | GCTTGATTTCCTCCTGCCTG |
| MGAT1 | 1 | ATCCTGGTCATCGCCTGTG | CATCCAGTCGTCCCAGAAGG |
| MGAT1 | 2 | GCTCTACCGCACCGACTTT | CCTAAGAGGGAAACACAGGC |

**Supplementary Table 3| Oligonucleotide sequences for amplifying the sgRNA target site in in avian (CLEC213 and DF-1) or human (A549) cell lines to confirm KO of targeted gene.**
